# Antisense oligonucleotide-mediated perturbation of long non-coding RNA reveals functional features in stem cells and across cell types

**DOI:** 10.1101/2022.02.16.480297

**Authors:** Chi Wai Yip, Chung-Chau Hon, Kayoko Yasuzawa, Divya M. Sivaraman, Jordan A. Ramilowski, Youtaro Shibayama, Saumya Agrawal, Anika V. Prabhu, Callum Parr, Jessica Severin, Yan Jun Lan, Josée Dostie, Andreas Petri, Hiromi Nishiyori-Sueki, Michihira Tagami, Masayoshi Itoh, Fernando López-Redondo, Tsukasa Kouno, Jen-Chien Chang, Joachim Luginbühl, Masaki Kato, Mitsuyoshi Murata, Wing Hin Yip, Xufeng Shu, Imad Abugessaisa, Akira Hasegawa, Harukazu Suzuki, Sakari Kauppinen, Ken Yagi, Yasushi Okazaki, Takeya Kasukawa, Michiel de Hoon, Piero Carninci, Jay W. Shin

## Abstract

Within the scope of FANTOM6 consortium, we perform a large-scale knockdown of 200 long non-coding RNAs (lncRNAs) in human induced pluripotent stem (iPS) cells, and systematically characterize their roles in self-renewal and pluripotency. We find 36 lncRNAs (18%) exhibiting cell growth inhibition From the knockdown of 123 lncRNAs with transcriptome profiling, 36 lncRNAs (29.3%) show molecular phenotypes. Integrating the molecular phenotypes with chromatin-interaction assays further reveals *cis*- and *trans*-interacting partners as potential primary targets. Additionally, cell type enrichment analysis identifies lncRNAs associated with pluripotency while the knockdown of *LINC02595*, *CATG00000090305.1* and *RP11-148B6.2* modulates colony formation of iPS cells. We compare our results with previously published fibroblasts phenotyping data and find that 2.9% of the lncRNAs exhibit consistent cell growth phenotype, whereas we observe 58.3% agreement in molecular phenotypes. This highlights molecular phenotyping is more comprehensive in revealing affected pathways.

## Introduction

The effort to characterize the human transcriptome^1, 2^ has revealed a large collection of long noncoding RNAs (lncRNAs), which are pervasively transcribed with little or no protein coding potential. Most lncRNAs are transcribed by RNA polymerase II and they are frequently capped, spliced and polyadenylated.^3^ Although an increasing number of lncRNAs have been identified in mediating multiple regulatory processes,^4^ experimentally curated functional lncRNAs represent less than 1% of all known lncRNA transcripts.^5, 6^ While most lncRNAs are expected to acquire function over evolutionary time,^7^ a genome wide analysis of 1,829 samples of human cell types or tissues further indicated 69% of lncRNAs with functional insights.^1^ To better address cell type specific functional lncRNAs, a loss-of-function genetic screen of lncRNAs is vital.^8^

Human induced pluripotent stem (iPS) cells are reprogrammed from somatic cells by introducing exogenous expression of the transcription factors OCT3/4, SOX2, KLF4 and c-Myc.^9^ Although minor differences amongst iPS cells^10^ and between iPS cells and embryonic stem (ES) cells^11^ are observed, iPS cells functionally resemble ES cells by forming embryoid bodies or teratomas, and by undergoing lineage specific differentiations. Recent studies in human ES cells and iPS cells showed indispensable roles of lncRNAs for their self-renewal and differentiation. For example, *lncRNA-ES1*, *lncRNA-ES3* and *FAST* play a role in maintaining pluripotency,^12, 13^ *LINC-ROR* in cell reprogramming^14^ and self-renewal,^15^ and *TUNA* in neuronal differentiation.^16^ Furthermore, pervasively expressed lncRNAs are associated with more active and open chromatin observed in pluripotent cells.^3^ LncRNAs with higher diversity and expression level provided pluripotent cells an attractive model in lncRNA functional annotation. Indeed, genetic screens of lncRNAs revealed iPS cells with the highest percentage of functional lncRNAs among several cell types.^17^

Genetic screens using DNA-targeting technologies such as CRISPRi^17^ and CRISPRa^18^ or RNA-targeting technologies such as RNAi^19^ and LNA gapmer antisense oligonucleotides (ASOs)^8^ have demonstrated systematic strategies to annotate novel lncRNA function. Many screening designs rely on the cellular phenotypic outcome, leading to the discovery of cell survival-related functions, while lncRNAs with other functional roles were less explored. In addition, due to the tissue-specific expression pattern of lncRNAs,^20^ the functions of specific sets of lncRNAs are only revealed under specific cell types and conditions. Studying the same lncRNAs across cell types and conditions will bring further insights into the lncRNA biology.

We have previously characterized 285 lncRNAs in human dermal fibroblast (HDF) primary cells by targeting the RNA molecules.^8^ In this study, we adopted LNA ASOs to directly target RNA molecules and streamlined this with cellular phenotyping and transcriptomic profiling to reveal functional roles. While the cell growth phenotype revealed lncRNA function in self-renewability of iPS cells, molecular phenotyping explored their roles in molecular pathways related to pluripotency and others. By perturbing 390 lncRNAs in the human iPS cells, we identified functional lncRNAs at both the cellular and molecular phenotypic levels (**Fig. 1**). RNA-DNA and DNA-DNA interaction data were integrated to the knockdown profiles to identify primary gene targets. Three lncRNAs regulating pluripotency were identified and validated. To comprehensively compare the properties and functions of lncRNAs across different cell types, we selected the lncRNAs robustly expressed in both HDF and iPS cells, and performed knockdown by the same set of ASOs. We observed cell type specificity in the growth phenotype but broader consistency in molecular phenotype, implying that distinct growth phenotypes may be due to different secondary responses between the cell types.

**Figure 1.**
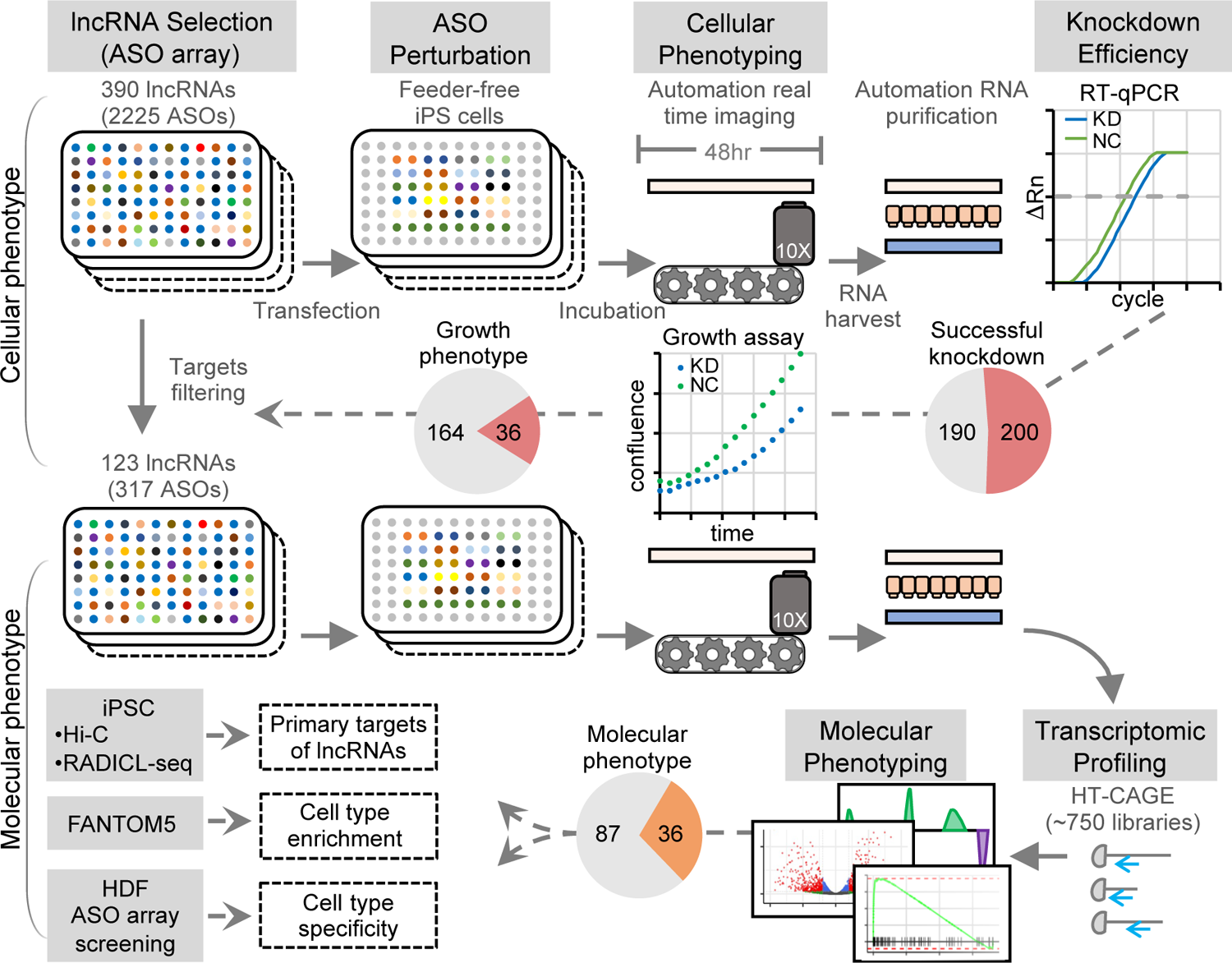
Schematic of the ASO array screen for growth phenotype and molecular phenotype of lncRNAs.

## Results

### Properties of 390 lncRNA targets

We selected 390 lncRNAs expressed in iPS cells and designed 5 to 10 ASOs per lncRNA to specifically suppress their expression in iPS cells. These lncRNAs cover a broad range of expression levels in iPS cells (**Fig. 2A**) and a broad range of cell type specificity amongst 71 cell types (**Fig. 2B**), with 102 lncRNAs overlapping with the targets tested in our previous study in HDF.^8^ These lncRNAs are generally categorized as either antisense, divergent, intergenic or sense intronic, and 12.3% of the lncRNAs are derived from enhancer-like regions (**Fig. 2C**). Subcellular fractionation analysis revealed 33% of the lncRNAs are enriched in the chromatin, 23% in the nucleoplasm and 44% in the cytoplasm (**Fig. 2D-E**). Notably, they were selected to represent lncRNAs commonly expressed in stem cells and their expression levels highly correlate with other iPS and ES cell lines (**Fig. 2F**).

**Figure 2.**
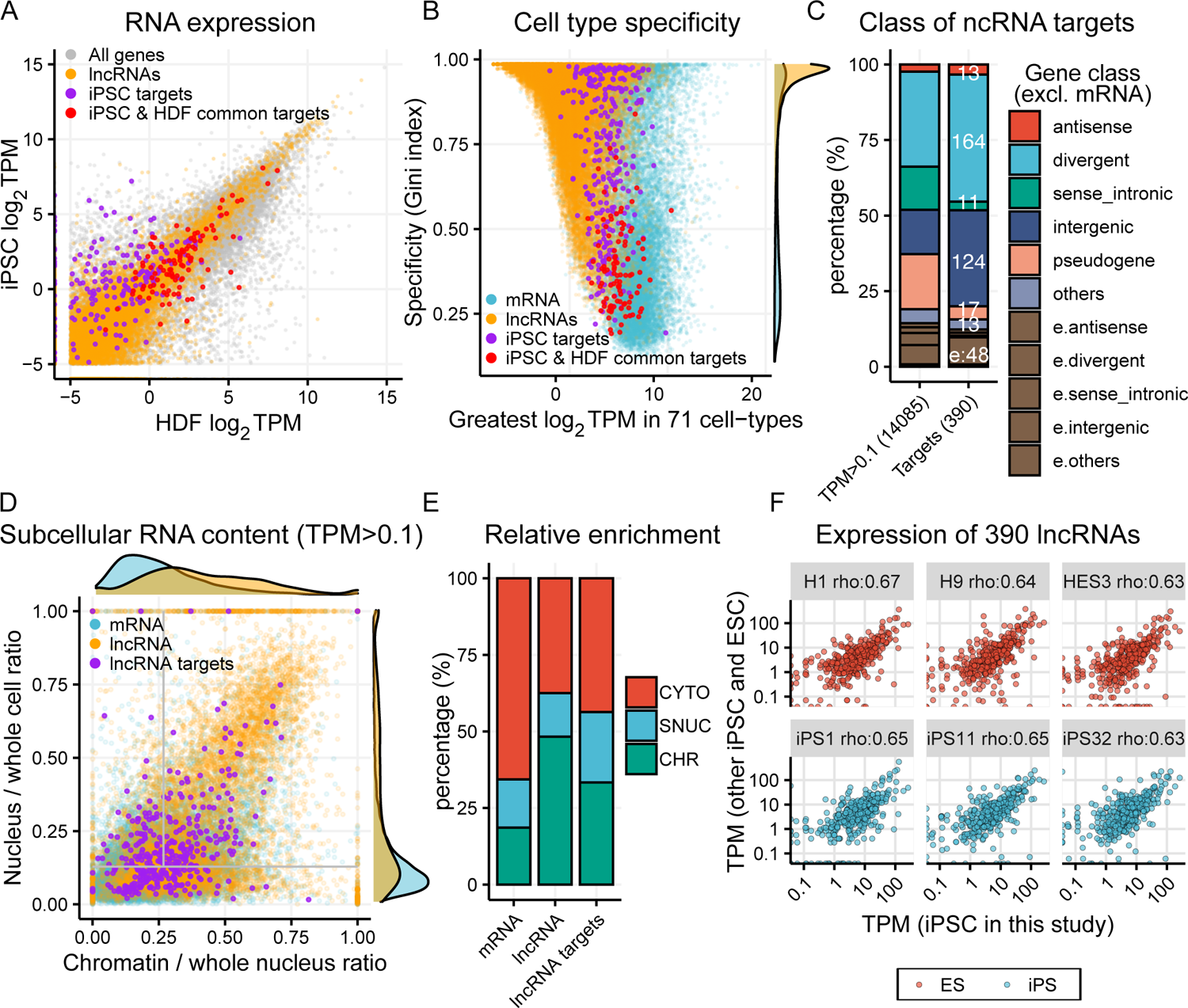
Properties of the 390 lncRNA targets. **A**, Gene expression level of iPSC and HDF derived from bulk RNA-sequencing on the native cells, highlighting the lncRNAs (orange), the screening targets in iPSC alone (purple) and the targets have been screened in both iPSC and HDF (red). **B**, Cell type specificity according to the expression of each gene in 71 cell types including iPS and ES. Gini index was calculated by comparing CAGE expression profiles of these cell types. **C**, Gene class of the 390 lncRNA targets and their corresponding proportion in iPSC when applying different cutoffs of expression level determined by bulk RNA-sequencing. Coding mRNA was excluded from the proportion. **D**, Subcellular localization of expressed genes (TPM > 0.1, determined by bulk-RNA seq) in iPSC, showing coding mRNA (blue), lncRNA (yellow) and the 387 targets (purple, 3 targets were undetermined). Subcellular fractionation of iPSC was performed and followed by RNA-sequencing on the cytoplasmic, soluble nucleoplasmic and chromatin fractions. The nuclear ratio from the whole cell (y-axis) was calculated from the normalized TPMs for (SNUC + CHR) / (CYTO + SNUC + CHR). The chromatin ratio from the whole nucleus (x-axis) was calculated from the normalized TPMs for CHR / (SNUC + CHR). The regions represent enrichment to CHR (upper-right), SNUC (upper-left) and CYTO (bottom) are segmented by the grey lines, which are the median value of all the genes (>1 TPM) of the 2 axes. **E**, The relative enrichment of mRNAs, lncRNAs and the lncRNA targets in the 3 subcellular compartments, defined by the criteria described in (**D**). **F** Expression level of the 390 lncRNAs in our iPS line and comparison to 3 human ES lines and 3 other human iPS lines (iPS1: hiPS; iPS11: iPSC11-CRL2429; iPSC32: iPSC32-CRL1502) from the FANTOM5 data.

### Cellular phenotyping unveils lncRNAs influencing self-renewal

After knocking down 390 lncRNAs with the ASOs (n = 2,225), we measured knockdown efficiency using RT-qPCR, and their effect on cell growth (i.e. cellular phenotype) using a real-time imaging-based growth assay, comparing against the non-targeting negative control (NC) ASO as background. We found 750 out of 2,225 ASOs with knockdown efficiency ≥ 50% (i.e. effective ASOs) and 200 lncRNAs with ≥ 2 effective ASOs (**Fig. 3A, Table S1**). The cell growth assay revealed 176 out of the 750 effective ASOs to significantly inhibit cell growth (Student’s t-test, *p*-value < 0.05; **Fig. 3B, S1A-B**). This observation was further supported by an independent optical density-based WST-1 growth assay, which strongly correlated with the image-based growth assay (**Fig. 3C**). Notably, ASOs with effective knockdown showed significantly stronger growth inhibition (Wilcoxon test, *p* < 0.05, **Fig. 3D**) and ASOs targeting the same lncRNA showed a significant concordance of the normalized growth rate (Methods; **Fig. 3E, S1C**). These results suggest that the observed growth inhibition is likely attributed to the specific effects of lncRNA knockdown instead of unspecific effects (e.g. ASO toxicity). Among the 200 lncRNAs with successful knockdown, 36 (∼18%) had 2 or more growth inhibiting ASOs (robust cutoff) while 11 (∼5.5%) were statistically significant based on binomial test at p-value < 0.05 (stringent cutoff; methods; **Fig. 3F**).

**Figure 3.**
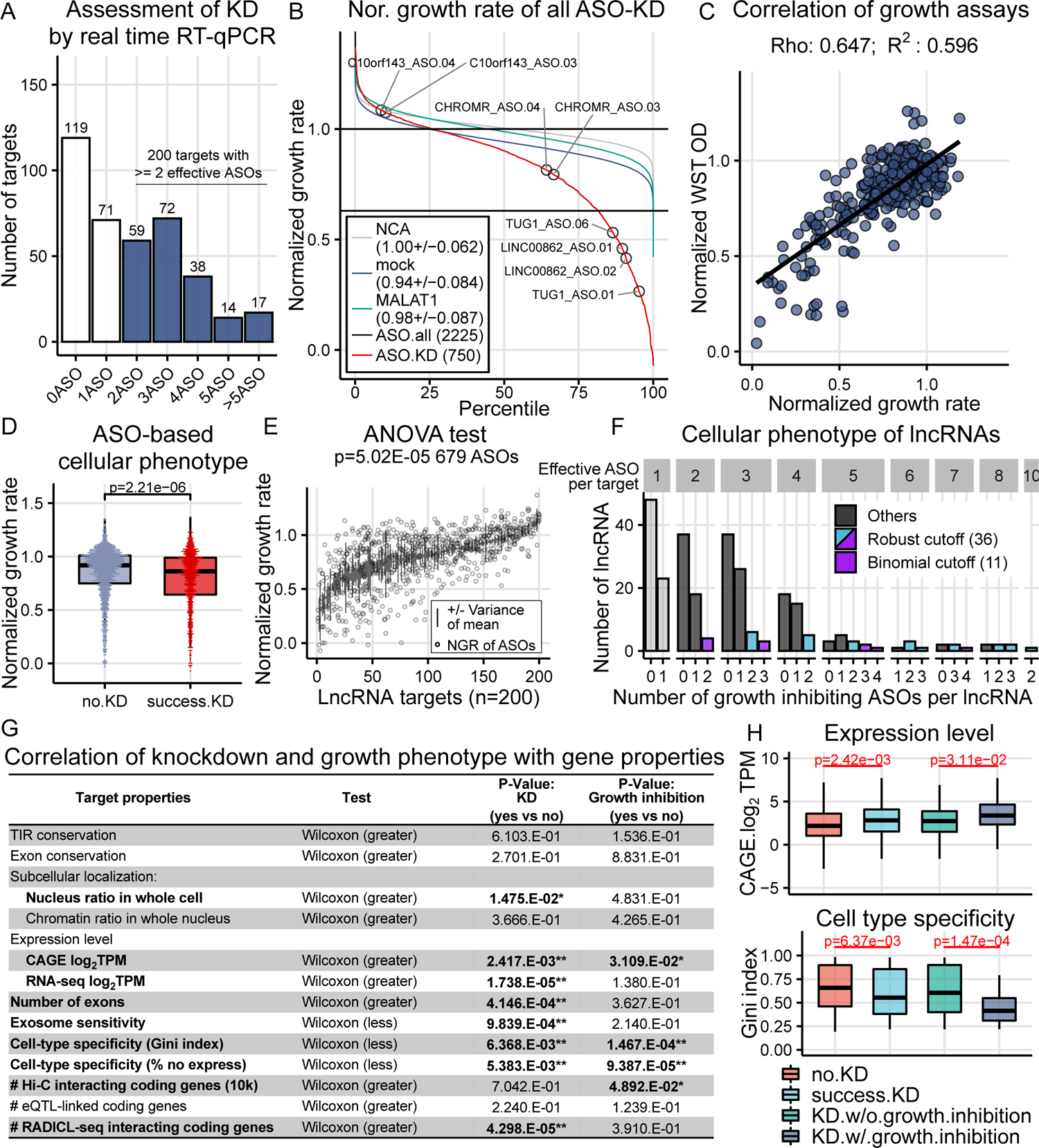
Cellular phenotypes and their correlations with genomic properties of lncRNAs. **A**, Number of target with successful KD and the number of ASO showed KD from the targets. Real time RT-qPCR was performed with 2 biological replicates. **B,** The distributions of normalized growth rate for all ASO KD. A cutoff of 0.63, which is 6 SD from the mean of the negative controls (cell transfected with NC ASO), was used to determine change of growth. **C,** Correlation between WST-1 growth assay and Incucyte growth assay in the ASO subset selected for CAGE profiling. Two biological replicates were included. **D,** Normalized growth rate of ASO with or without showing successful knockdown. **E**, Normalized growth rate of each ASO targeting the same lncRNAs and the variances of each lncRNA (679 ASOs for 200 lncRNAs). One-way ANOVA test resulted in a *p*-value as 5.02E-5. **F,** LncRNAs are defined as growth phenotype by at least 2 effective ASOs showing growth inhibition (robust cutoff, n= 36) and by the number of ASO passing binomial statistics (stringent cutoff, n=11). **G**, Table showing the *p*-value of the correlation tests between the genomic properties and the result of KD and growth phenotype (lncRNA targets, n=390; no.KD, n=190; KD, n=200; KD.growth, n=36; KD.no.growth, n=164; * <0.05, ** <0.01). **H**, Wilcoxon tests on expression level and cell type specificity.

We next asked whether different properties of the lncRNAs correlate with the successful knockdown (≥ 2 effective ASOs) and growth inhibition (≥ 2 growth inhibiting ASOs). We found that lncRNAs with successful knockdown are characterized by more enriched in the nucleus, higher number of exons, higher expression levels, lower exosome sensitivity, and lower cell type specificity (**Fig. 3G, S1D**). Additionally, lncRNAs with growth inhibition exhibited significantly higher expression level and lower cell type specificity (**Fig. 3G-H, Table S2**).

### Molecular phenotypes recapitulate cellular phenotypes

To investigate the effect of lncRNA knockdown on gene expression profile (i.e. molecular phenotype), we performed Cap Analysis of Gene Expression (CAGE)^21, 22^ after knockdown with selected effective ASOs (n = 317; targeting 123 lncRNAs). For each ASO, we performed differential gene expression, pathway enrichment and differential transcription factor (TF) binding motif analyses, comparing against the NC ASO background (Methods). First, to identify lncRNAs whose knockdown led to reproducible and significant transcriptome global changes, we paired the ASOs targeting the same lncRNA (same-target-ASO-pair; 293 pairs) and compared the mean distance of each pair against a null distribution of permuted pairs of NC ASO replicates (NC-ASO-pairs). We quantified the distance based on the overall extent of transcriptome changes for each ASO using Kolmogorov’s *D* (refer to as distance; Methods). This resulted in 52 significant same-target-ASO-pairs targeting 28 (∼22.7%) lncRNAs (empirical *p*-value < 0.01 and Wilcoxon test FDR < 0.05; Methods; **Fig. 4A**). We observed a strong correlation between the overall extent of transcriptome changes and growth inhibition (**Fig. S2A-B**), suggesting that gene programs such as apoptosis or inhibition of cell growth induces global shift in the molecular phenotype. Consistently, pathways related to cell growth, e.g. cell cycle and cell death, are highly correlated with the normalized growth rate (Methods, **Fig. 4B, S2C**). These analyses revealed a consistent agreement between two independent measurements: the cellular and molecular phenotypes.

**Figure 4.**
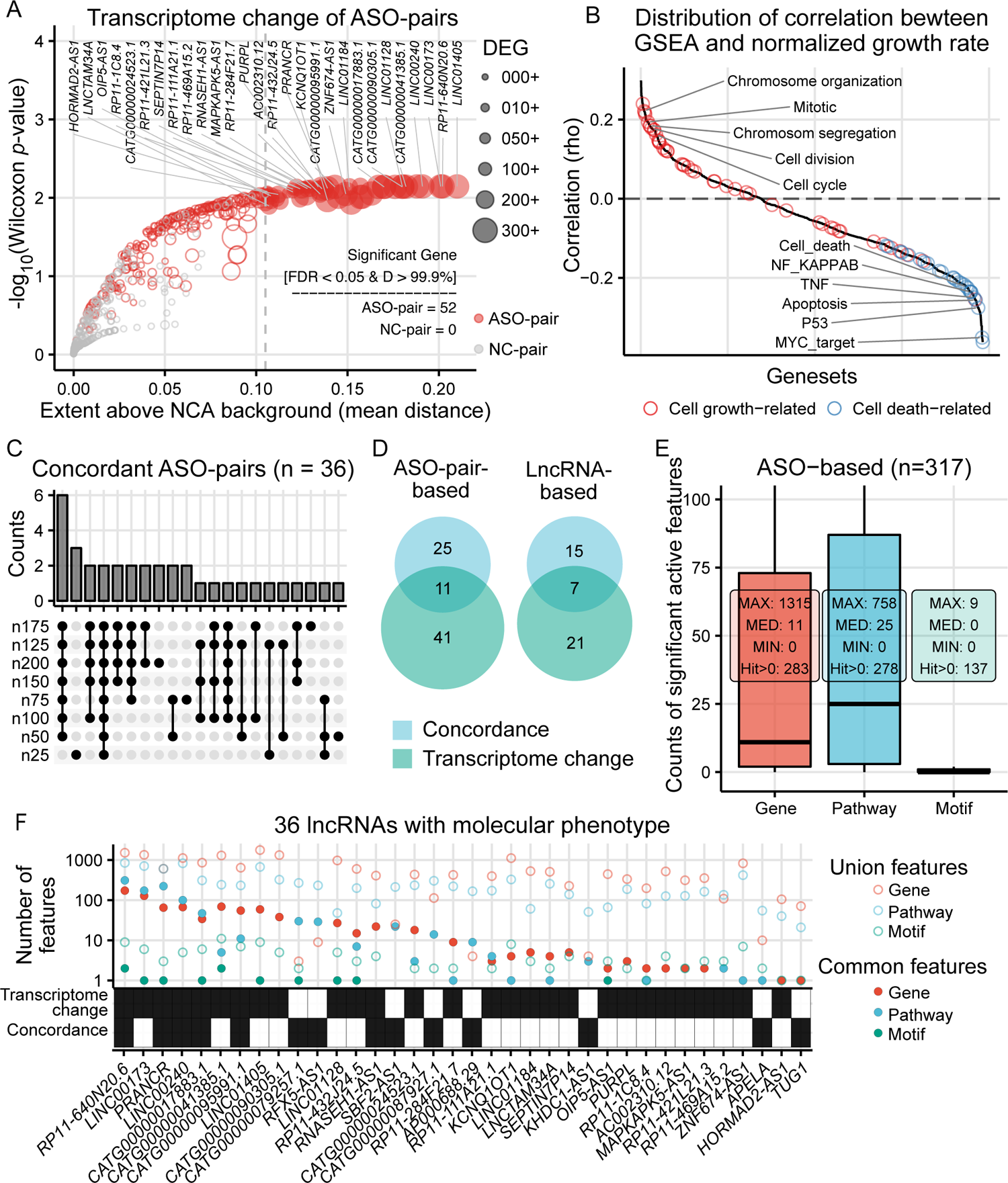
Molecular phenotypes of 123 lncRNAs. **A**, Visualization of the transcriptome changes of ASO-pairs after edgeR analysis. 52 ASO-pairs were considered to have significant transcriptomic change. **B**, Correlation between GSEA result for each pathway with the normalized growth rate across all the ASOs. **C**, Concordant ASO-pairs derived from different geneset sizes. **D**, ASO-pair-based and lncRNA-based Venn diagram showing the overlap of pairs/lncRNAs that are concordant and with reproducible transcriptome change. **E**, ASO-based results of number of significant differentially active molecular features in both up- and down-regulations (gene: *edgeR* FDR < 0.1, |across sample Z-Score| > 1.645; pathway: *fgsea* padj < 0.05, |across sample Z-Score| > 1.645; motif: MARA FDR < 0.1. **F**, Number of union significant features and common significant features of representative ASO-pairs of 36 lncRNAs showing molecular phenotype. The target lncRNAs are excluded from the number of differentially active genes. The representative ASO-pairs and the contents of significant features are shown in **Table S3**.

### Analysis of molecular phenotype indicates functional aspect of lncRNAs

To investigate molecular features impacted by lncRNA knockdown, we first implemented the Jaccard index to test the concordance of differentially expressed genes (DEGs) (Methods). We found that the mean Jaccard index of the same-target-ASO-pairs is significantly higher than the null distribution from permuted different-target-ASO-pairs (empirical *p*-value < 0.05; **Fig. S2D**), suggesting that the observed ASO effects are target-specific. This observation is also supported by a more stringent analysis incorporating the Wilcoxon test (empirical FDR < 0.05; **Fig. S2E-F**).

To identify lncRNAs with reproducible molecular phenotypes from 2 or more ASOs, we implemented a reciprocal enrichment method to reveal ASO-pairs leading to concordant transcriptomic effects (Methods; **Fig. S3A-C**). This method significantly distinguishes same-target-ASO-pairs against different-target-ASO-pairs (empirical *p*-value < 0.05; **Fig. S3D**). Overall, 36 ASO-pairs (**Fig. 4C, S3E**), which account for 22 lncRNAs, showed concordant transcriptomic effects upon knockdown. Hereby, we identified 43 lncRNAs which showed reproducible transcriptome global changes (n = 28; **Fig. 4A**) and 22 lncRNAs with concordant transcriptomic effects with the support of ≥ 2 ASOs (**Fig. 4D**).

Next, we sought to define the optimal cutoffs for identifying significant molecular features (i.e. genes, pathways and TF binding motifs) after ASO knockdown, by maximizing the ratio of concordance of same-target-ASO-pairs to different-target-ASO-pairs (Methods; **Fig. S3F**). At these optimal cutoffs, the median number of significant genes, pathways and motifs of the 317 ASO knockdowns were respectively, 11, 25 and 0 (**Fig. 4E**). Among the 43 lncRNAs with reproducible knockdown effects, 36 of them showed at least one common significant molecular feature. These 36 lncRNAs are defined as showing functionally reproducible molecular phenotypes (**Fig. 4F, Table S3**). We compared these 36 lncRNAs with the remaining targets (n = 87) for genomic property enrichment analyses but no significant enrichment was found (**Table S2**). Among these 36 lncRNAs, only 59 out of 111 same-target-ASO-pairs are concordant. To explore the cause of discordant ASO-pairs found in these 36 lncRNAs, we compared isoform-targets of the two ASOs in a pair and showed that ASOs from the discordant pairs are significantly more different in isoform coverage (*p*-value = 0.0214; Methods; **Fig. S3G**).

### Primary gene target at chromatin level

LncRNAs regulate cellular activity through multiple mechanisms,^23^ while the majority of the experimentally curated lncRNAs act via transcriptional regulation.^6^ However, the primary gene target of lncRNA is not distinguishable from the secondary responses in the molecular phenotype. To predict the primary effects of lncRNAs from their physical associations with chromatin, we integrated the molecular phenotype with Hi-C and RADICL-seq, a method to capture RNA-DNA interactions,^24^ from the native iPS cells.

This section focused on the ASOs derived from the 36 concordant lncRNAs in molecular phenotyping while the results of other ASOs are listed in **Table S4**. We identified the protein-coding genes in contact with lncRNAs either from their DNA loci or the RNA molecules, which are reflected by Hi-C and RADICL-seq respectively. From the 36 lncRNAs assayed for transcriptomic response, 35 lncRNA loci showed significant Hi-C contacts (FDR < 0.05), resulted in 995 lncRNA-protein coding gene pairs. From the RADICL-seq results, 17 lncRNAs showed significant contacts (FDR < 0.05) and 661 lncRNA-protein coding gene pairs were identified. Consistent with previous studies,^25, 26^ the significant pairs of DNA-DNA and RNA-DNA interactions were largely overlapped (**Fig. S4A**). We thereby considered the 995 pairs with Hi-C contacts as *cis* interactions (35 lncRNAs) while those pairs specific to RADICL-seq, with at least 1 Mb apart, as *trans* interactions (n = 270, from 17 lncRNAs). Additionally, eQTL-genes^27^ with the variants located in the promoters and gene bodies of the lncRNA targets were integrated, showing 31 lncRNAs with detectable eQTL-genes (**Fig. S4B**).

By incorporating the significant DEGs with significant contact genes of the lncRNAs, we showed that *cis* and *trans* interactions are globally enriched with DEGs as compared to background (Methods; **Fig. S4C**). Then, we linked the significant DEGs with *cis* interactions for the concordant lncRNAs and identified 18 of them with potential primary targets in *cis* (**Fig. 5A**, full list including non-concordant ASOs in **Table S4**). One of the significant pairs is derived from *TUG1* ASO without global concordance. Its knockdown mediates down-regulation of a protein-coding gene *SELENOM*, which is located in the same TAD, with significant Hi-C and RADICL-seq contacts from the *TUG1* locus (**Fig. 5B**). We identified another ASO causing similar down-regulation of *SELENOM* in CAGE and independently validated it by RT-qPCR (**Fig. 5C**). *TUG1* and *SELENOM* share similar phenotypic outcomes where they mediate cell proliferation in multiple cancers^28–31^ and obesity.^32, 33^ In addition, we revealed that the lncRNA *RNASEH1-AS1*, without affecting the expression of *RNASEH1*, regulated *ADI1* in *cis*, supported by an independent knockdown and RT-qPCR (**Fig. S4D-E**).

**Figure 5.**
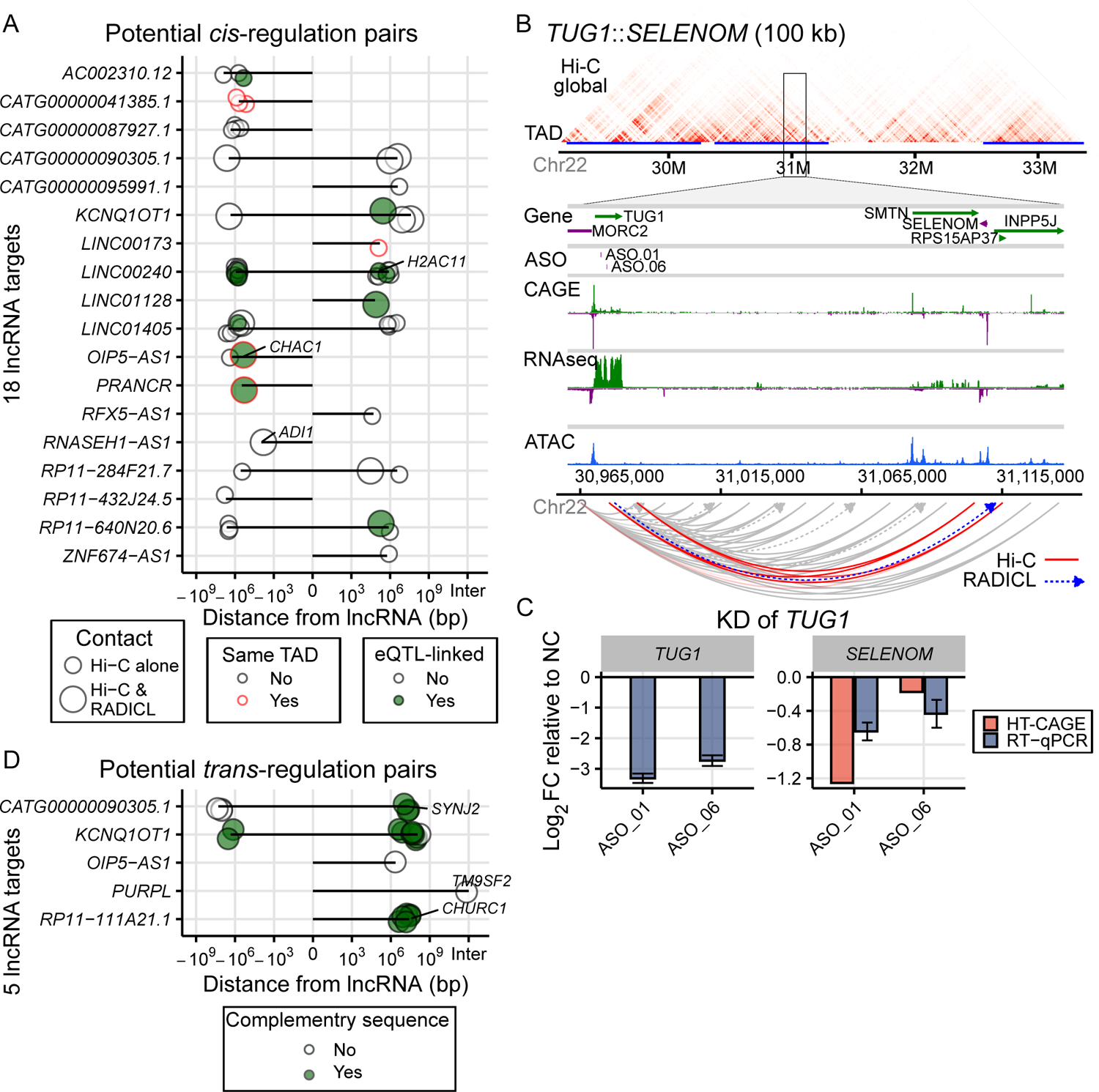
Potential primary target of lncRNAs. **A**, A list of potential *cis*-regulation lncRNAs and their primary protein coding gene targets which are regulated significantly after knockdown (FDR < 0.1 & |Zscore| > 1.645). Hi-C contacts and TAD regions were defined at the resolutions of 10 kb and 50 kb, respectively. Within TAD: both contact regions located in the same TAD; eQTL-linked: linked-pairs with FDR < 0.25 from any of the 49 tissue samples. **B**, Genomic view of the *cis*-interaction between *TUG1* and *SELENOM*. **C**, Down-regulation of *SELENOM* upon ASO KD of *TUG1* from CAGE and an independent knockdown and RT-qPCR. Two biological replicates were included for each ASO. **D**, Five lncRNAs with potential *trans*-regulation for having *trans*-interacting protein coding genes (>1 Mb distance, no significant Hi-C contacts, detectable in HT-CAGE) with differential expression after knockdown (FDR < 0.1 & |Zscore| > 1.645). Complementary sequence similarity found from RNA transcripts towards the gene body of the protein coding genes.

Next, we performed an enrichment test on the *cis*-contacted protein-coding genes to the ranked DEGs for each lncRNA (Methods). The result revealed a significant enrichment of *cis*-contacts for *CATG00000087927* and *LINC00240* (adjusted *p*-value < 0.05; **Fig. S4F**), implying that these lncRNAs may exhibit strong *cis*-regulatory effects towards multiple genes spatially proximal and regulate cell growth pathways (**Fig. S4G**). Enrichment analysis on the eQTL-genes with the variants located in the lncRNA targets further highlighted *LINC00240*, at which multiple eQTL-genes were regulated upon the knockdown of *LINC00240* (**Fig. S4H**).

We integrated the transcriptomic profiles with *trans* interactions to investigate potential *trans*-acting lncRNAs. We performed pair-base identification combined with complementary sequence similarity detection between lncRNA targets and gene DNA sequences, and discovered 5 potential *trans*-acting lncRNAs (**Fig. 5D**, full list in **Table S4**) including *CATG00000090305* (**Fig. S5A-B**). Enrichment analysis, which reveals multiple genes regulation did not show significant (**Fig. S5C**). Overall, we showed that the integration of the genome-wide static chromatin interaction data together with large-scale loss-of-function transcriptomic responses predict primary target genes of lncRNAs.

### LncRNAs modulate cell identities

To elucidate the differentiation potential of lncRNA in pluripotent stem cells, we tested whether lncRNA knockdown leads to gain or loss of predefined cell type molecular signatures. We derived the cell type specificity score from 71 cell types including iPS and ES cells by applying the Gini index (Methods; **Table S5**). After identifying the top specific genes from each cell type as genesets (Methods), we performed geneset enrichment analysis against the molecular phenotype data. We additionally included genesets from *POU5F1* knockdown transcriptomic responses to represent pluripotency-related genes.

Among the 36 concordant lncRNAs, we identified 16 lncRNAs with enrichment in various cell types (**Fig. 6A**). The knockdown profiles from *POU5F1* and 4 other lncRNAs showed significant enrichment of stem cell related genesets. From the 87 lncRNAs lacking global concordance, we found 24 of them showing cell type enrichment with the support of at least 2 ASOs (**Table S5**). The results of *POU5F1* knockdown and some known lncRNAs demonstrated the applicability of this analysis. For example, knockdown of *NUTM2B-AS1*, which was recently shown to be involved in a neuromuscular disorder,^34^ was enriched with neurons. Knockdown of *TUG1* reflected negative enrichment of osteoblast, tendon and fat cells, with *TUG1* promoting osteogenic differentiation from periodontal ligament stem cells^35^ and tendon stem cells,^36^ as well as promoting brown remodeling of white adipose tissue.^33^

**Figure 6.**
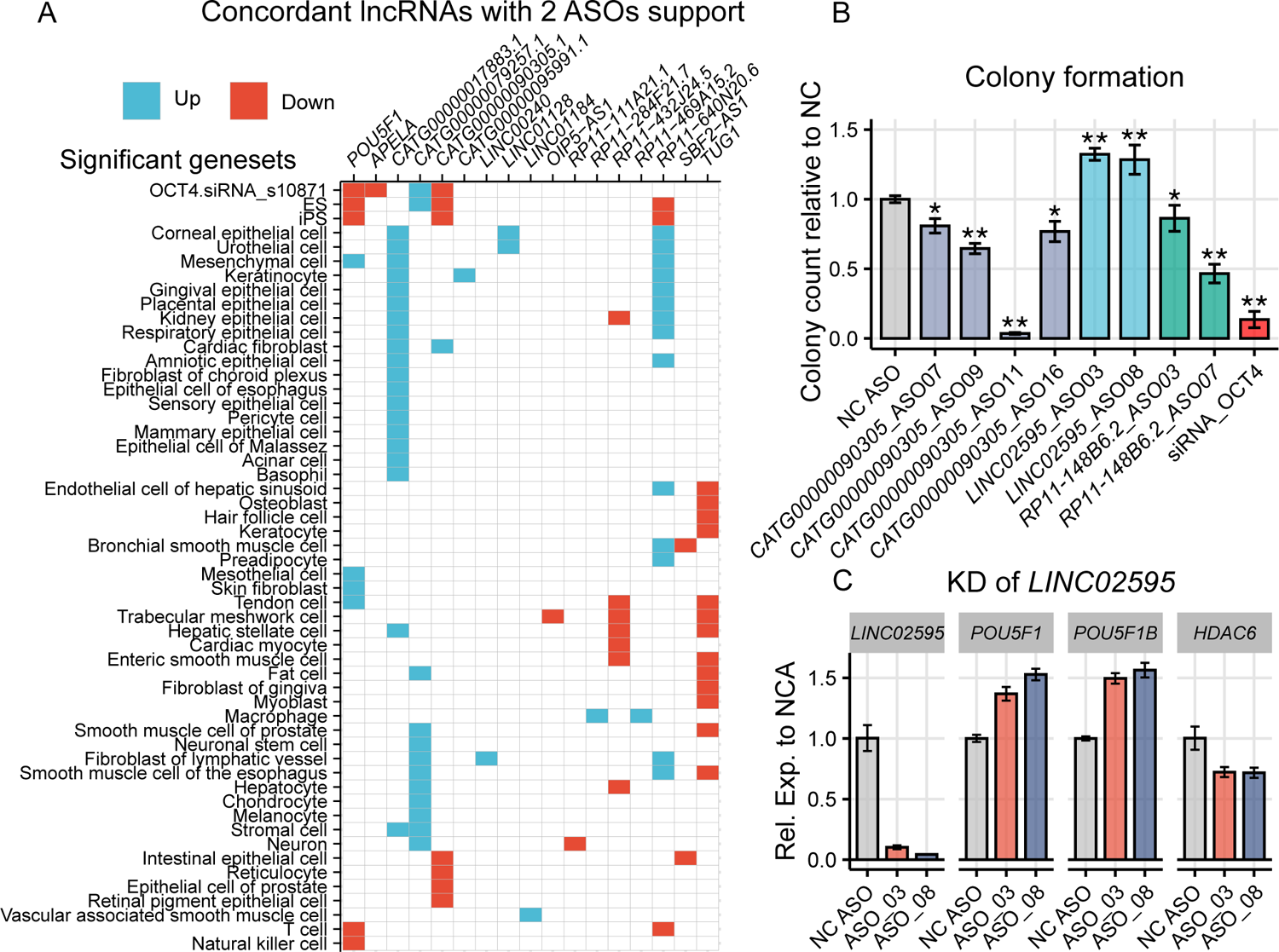
Enrichment of cell type specific genes upon lncRNA perturbation. **A**, Expressed genes from a cell type were ranked according to the specificity scores derived from Gini index of 71 cell types. Expression level of a gene in a cell type in the lower 5 percentile among the 71 cell types is considered non-expressed. Eight genesets were generated from each cell type by selecting the top 25, 50, 75, 100, 125, 150, 175 and 200 cell type specific genes. Genesets were tested for enrichment in all the ASO-KD profiles to acquire p.adj of *fgsea* and across sample Z-score on NES. LncRNA targets having at least 2 ASOs with NES in the same direction, p.adj < 0.05 and |Z-score| > 1.645 are shown in the heatmap. Only concordant lncRNAs and POU5F1 control are shown, full list is in **Table. S5**. **B**, Colony formation assay upon the knockdown of *CATG00000090305*, *LINC02595*, *RP11-148B6.2* and OCT4. The number of colonies were counted 6 days after the cells were seeded by crystal violet staining. The chart shows the mean and standard deviation of at least 3 biological replicates. **C**, Knockdown of *LINC02595* with spontaneous differentiation condition, RNA levels relative to NC ASO-transfected samples of *LINC02595*, *POU5F1*, *POU5F1B* and *HDAC6* are shown. Two biological replicates were included for ASO knockdown.

To functionally validate lncRNAs that modulate stem-cell identity, we performed colony formation assays with lncRNA knockdown. The knockdown of the pluripotency-supporting lncRNAs *CATG00000090305.1* and *RP11-148B6.2*, showed a decrease in colony formation. Alternatively, the knockdown of *LINC02595*, which disrupts stem cell molecular phenotype, increased the number of colonies (**Fig. 6B**). This demonstrates that geneset enrichment analysis predicts function of lncRNA and that these lncRNAs affect the cell survival of iPS cells at the single cell stage.

We further investigated the role of *LINC02595*, for its role in disrupting the stem cell molecular phenotype. Knockdown of *LINC02595* by 2 independent ASOs down-regulated *HDAC6* and up-regulated *POU5F1* and *POU5F1B* under spontaneous differentiation conditions (**Fig. 6C**). Notably, *HDAC6* was shown to be regulated by *LINC02595* in *trans* (**Table S4**) and *HDAC6* level increased along differentiation from iPS cells to neurons in our previous study (FANTOM5^1^). *HDAC6* encode for histone deacetylase and suppresses multiple genes during differentiation.^37^ Inhibition of it up-regulated OCT4 in porcine blastocysts^38^ while HDAC inhibitor was shown to improve iPS cell reprogramming from peripheral blood mononuclear cells.^39^

### Extent of lncRNA function across cell types

LncRNAs are known to exhibit cell type- and tissue-specific expression, and cell type specificity is generally higher than mRNA (**Fig. 7A**).^20^ To explore the role of distinct subcellular localization of lncRNAs in different cell types, we performed fractionation on HDF and iPS cells for subcellular RNA-seq and showed that subcellular enrichment is similar between lncRNAs and mRNAs across all the expression levels (**Fig. 7B**). Consistent with a previous study,^40^ this analysis suggests that many lncRNAs tend to maintain their subcellular localization, which largely dictates the functions of lncRNAs.^41^ Thus, we reasoned that lncRNAs that express ubiquitously across cell types exhibit similar molecular roles.

**Figure 7.**
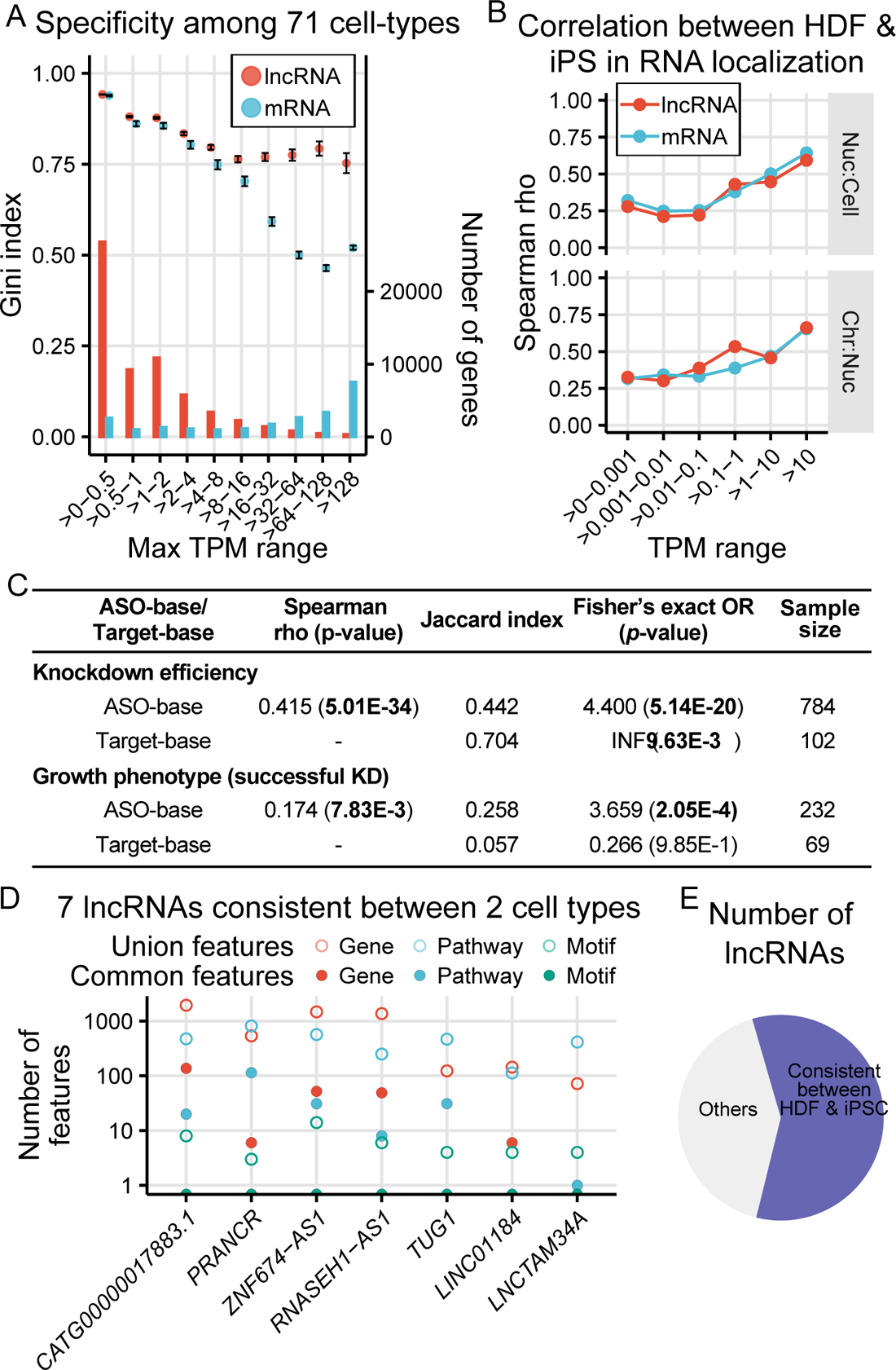
Comparison of lncRNAs between HDF and iPS cells. **A**, Cell type specificity at expression level. Gini index was calculated by comparing CAGE expression profiles of 71 cell types including iPS and ES cells. The mean Gini index and 95% confidence interval was shown for mRNA and lncRNA at each expression level bin, which was determined by the greatest TPM of each gene among the 71 cell types. **B**, Spearman correlation of the nucleus-to-whole cell ratio and the chromatin-to-whole nucleus ratio between HDF and iPS cells. The correlation was divided into mRNA and lncRNA in different expression level bins. **C**, The comparison of KD efficiency and growth phenotype of the same sets of lncRNA targets between HDF and iPS cells. For Jaccard index and Fisher’s exact, the cutoff of KD is 50% while the cut off of growth phenotype is normalized growth rate < 0.63. OR: odds ratio. **D**, Number of union significant features and common significant features of representative ASO-pairs of 7 lncRNAs showing consistent molecular phenotype across HDF and iPS cells. The target lncRNAs are excluded from the number of differentially active genes. The representative ASO-pairs and the contents of significant features are shown in **Table S6**. **E**, Pie chart summarizing the number of lncRNAs with consistent molecular phenotype across HDF and iPS cells.

To test this hypothesis, we analyzed lncRNAs from two functional genetic screens: HDF^8^ and iPS cells (this study). We found 102 lncRNA targets that are common and have comparable expression levels and subcellular localizations in both cell types, with a wide range of specificity scores (**Fig. 2A-B, S6A**). Among them, 69 lncRNAs showed successful knockdown by at least 2 independent ASOs in both cell types and exhibited a positive correlation for the knockdown efficiency between two cell types (**Fig. 7C, S6B-C**). However, the cell growth phenotype correlation was low (2 overlapping lncRNAs; Jaccard index of 5.7%; **Fig. 7C, S6D-E**), which agrees with a previous study for the distinct growth phenotypes mediated by the same lncRNA loci across cell types.^17^ Additionally, the differential expression level of lncRNAs between HDF and iPS cells was found to contribute to the distinct growth phenotypes (**Fig. S6F**), which echoes our finding that the expression level of lncRNAs correlate with growth phenotypes (**Fig. 3G-H**).

We took advantage of the molecular phenotype data of 33 common lncRNAs and explored the lncRNA functional specificity across the two cell types. Among them, 12 lncRNAs were defined as concordant according to the molecular phenotyping. We included all 33 lncRNAs to increase the size of background but we focused on the results only from the 12 concordant lncRNAs. The results of the remaining 21 lncRNAs are listed in **Table S6**. To compare molecular phenotypes from two cell types, we performed the reciprocal enrichment test for ASOs targeting the same lncRNA (Methods; **Fig. S3A-C**). Notably, the same-target-ASO-pairs across the two cell types resulted in a higher concordance than the different-target-ASO-pairs (empirical *p*-value < 0.0001; **Fig. S7A**). Among the 12 concordant lncRNAs, 18 ASO pairs, or 8 lncRNA targets, showed concordant transcriptomic effect across HDF and iPS cells (**Fig. S7B, Table S6**).

Next, we extracted the significant molecular features (genes, pathways and TF binding motifs) of the 12 lncRNAs from HDF and iPS cells (**Fig. S7C**). We showed that the same-target-ASO-pairs are globally concordant for differentially expressed genes and pathways compared to different-target background (**Fig. S7D**), however, no common TF binding motif was found under the current threshold. From the 8 concordant lncRNAs, we required at least one common molecular feature to define functionally reproducible molecular phenotype across HDF and iPS cells. This resulted in 16 ASO-pairs targeting 7 lncRNAs (**Fig. 7D-E**). Furthermore, the significant differential expressed genes and pathways among these 7 lncRNAs are mostly unique (**Fig. S7E**), which excludes the possibility of off-target stress effects contributed by the ASOs. Overall, we revealed that, for lncRNAs expressed in both cell types, they generally show consistent molecular features across cell types while the cellular phenotype differs. The difference between the two phenotyping suggests that molecular phenotyping is more sensitive in revealing affected pathways during perturbation as compared to singular measurement such as cell growth.

To highlight lncRNAs shared molecular features between HDF and iPS cells, we revealed *TUG1* in regulating *ASCC2*, *SPRY4-AS1* in regulating *ARHGAP26* and *KHDC1-AS1* in regulating *KCNQ5*, while all these pairs showed significant Hi-C contacts in both HDF and iPS cells and the observation is supported by two independent ASOs (**Fig. S7F-H**). In the case of the *KHDC1-AS1*, although we did not observe its global concordance between the two cell types, we found concordant *cis*-regulatory function when the same ASOs were compared. This may be due to different transcript isoforms being targeted by different ASOs, suggesting that the molecular concordant events may be under-estimated.

## Discussion

We devised a systematic workflow to functionally characterize 200 lncRNAs in human iPS cells through both cellular and molecular phenotyping upon ASO-mediated knockdown. The large-scale perturbation study has revealed a scalable and effective way to evaluate their functional roles and potential primary targets in iPS cells. Here, we annotated 36 lncRNAs affecting cell growth (**Fig. 3F**) and 36 lncRNAs that elicit robust molecular features (**Fig. 4F**), while the corresponding ASOs and lncRNAs are listed respectively in **Table S2** and **Table S3**. The molecular phenotyping was extended to identify of *cis*- and *trans*-acting lncRNAs and lncRNAs that affect stem cell identity.

We report that 18.3% of the lncRNAs exhibit cell growth phenotype. This rate is comparable to our previous ASO array screen on human dermal fibroblasts.^8^ Previous study based on a CRISPRi pool screening on human iPS cells showed that 5.9% of lncRNA loci affect cell growth.^17^ Comparing the 65 common lncRNAs between CRISPRi pool screening and our study, we found only five lncRNAs with growth phenotype in both studies (**Fig. S1E-F**). The inconsistency could be partly due to the perturbation methods whereby CRISPRi targets DNA regulatory elements that may function independently from the RNA.^42, 43^ On the contrary, ASOs could degrade the nascent RNAs, which affect their transcription activities^44^ and transcription regulators recruitment in *cis*, and/or degrade mature RNAs in *trans*. Other explanations could be due to the strategy of stable or transient suppression, variations in iPS cell clones, methods to measure cell growth, and the non-targeting controls used, etc.

By targeting the transcript, ASO showed enhanced knockdown efficiency for lncRNAs with nuclear enrichment and higher expression levels. It has been reported that ASOs mediate the degradation of nuclear RNA including nascent RNA effectively relying on the higher enrichment of endogenous RNase H1 in the nucleus.^45^ This supports the knockdown of lncRNAs which are largely found to regulate genes at the chromatin level in the nucleus.^6^ Moreover, highly expressed and universally expressed lncRNAs are more likely to affect cell growth (**Fig. 3H**). This suggests that a set of ubiquitously expressed lncRNAs is indispensable and mediates essential pathways such as cell cycle progression, even though lncRNAs have been previously considered cell type specific. It is possible that the roles of cell type specific lncRNAs may only be activated by specific stimulation such as differentiation.

Previous studies have highlighted that the same lncRNA expressed in two different cell types may exhibit distinct cellular functions.^17^ In this study, we observed cell type dependent response to cell growth phenotype when the same lncRNAs were suppressed in two cell types.. However, we broadly observed a high degree of agreement in the molecular phenotype across both cell types (**Fig. 7E**), suggesting that the same molecular function of a lncRNA may affect the cell growth phenotype differently in different cell types. However, the concordant growth phenotype is generally underestimated due to the varying knockdown efficiency, which may not affect molecular phenotyping to the same degree when multiple genes from the transcriptome are analyzed. Therefore, relying solely on cell growth assays significantly underestimates functional classification of lncRNAs.

Integration analyses linked chromatin interaction assays and the knockdown data to identify the potential primary targets of lncRNAs. The enrichment analyses highlighted lncRNAs that affect the expression of multiple genes nearby in *cis*. For instance, *LINC00240*, which was shown to promote cancer cell growth,^46^ supports the expression of several histone genes in the nearby locus with significant Hi-C contacts and eQTL linkages (**Fig. 5A**). Additionally, we highlighted the *cis*-regulatory role of *TUG1* in controlling *SELENOM*. This *cis*-regulation was also found in mouse after *Tug1* knockout,^47^ however, the direction of regulation of the *cis*-genes including *SELENOM* (*Selm*) and *MORC2* (*Morc2a*) are opposite to our findings (**Table S4**). In addition to species-specific lncRNA function, the distinct *cis*-responses may be caused by the different methods of perturbation. While the exact dynamics to evaluate the cause-and-effect of lncRNA knockdown is still predictive, the integrative analysis provides an in-depth insight into potential regulatory processes for downstream analysis. A study including multiple time points will be useful for dissecting the precise dynamic mechanisms of lncRNAs in the future.

In summary, our study delineates a robust high-throughput lncRNA-perturbation, real-time imaging and transcriptomic profiling in a systematic fashion. This combined strategy reveals lncRNAs with functional annotation by cellular and molecular phenotyping in iPS cells. Going forward, systematic gene perturbation in multiple cell lineages together with additional multi-omics profiling will be necessary to fully annotate the human genome.

### Limitations of the study

This study is presented as a functional genomics screen with potential false negatives. LncRNA targets without reproducible growth phenotype or molecular phenotype may be due to insufficient sensitivity of the assays. In addition, this study is limited by the off-target effects of ASOs where responses unrelated to the target knockdown may interfere with phenotyping. To overcome potential off-target effects in the study, we introduced stringent criteria for molecular phenotyping, from which we reported a list of 36 lncRNAs with at least 2 concordant ASOs. While ASOs were used to directly target the RNA molecules, it is possible that ASO can also interfere with the transcription process when it targets the nascent transcript.^44, 48^ Therefore, *cis*-regulatory effect of lncRNA may be driven either by the lncRNA itself or the act of transcription. Finally, the scale of our study is limited to the number of assayed lncRNAs: 200 for cellular phenotyping and 123 for molecular phenotyping. For cell type specificity between HDF and iPS cells, we profiled 69 for cellular phenotyping and 12 for molecular phenotyping. This scale may affect the broader implications of biological nature of lncRNAs in the global analyses (shown in **Fig. 3G** & **Fig. 7C-E**).

## Acknowledgement

We thank Dr. Daisuke Yamada and Dr. Haruhiko Koseki for the generation of the iPS cell line. We also thank the Laboratory for Comprehensive Genomic Analysis in RIKEN for performing the CAGE library preparation and the next generation sequencing in this study. FANTOM6 was made possible by a Research Grant for RIKEN Center for Life Science Technology, Division of Genomic Technologies (CLST DGT) and RIKEN Center for Integrative Medical Sciences (IMS) from MEXT, Japan.

## Author’s contribution

Conceptualization: C.W.Y., J.W.S.; Investigation: C.W.Y., C.H., K. Yasuzawa, D.M.S., J.A.R., Y.S., S.A, A.V.P., C.P.; Experimental analysis: C.W.Y., K. Yasuzawa, D.M.S., Y.S., A.V.P., C.P., Y.J.L., F.L., T.K., J.D., H.N., M.I., J.L., M.K., M.M., W.H.Y.; Computational analysis: C.W.Y., C.H., J.A.R., S.A, A.P., J.C., M.T., M.d.H., X.S.; Database management: I.A., A.H., T.K.; Genome browser visualization: J.S.; Writing (original draft): C.W.Y., J.W.S.; Writing (review & editing): C.H., D.M.S., J.A.R., Y.S., S.A, A.V.P., C.P., M.d.H., P.C.; Project supervision: C.W.Y., C.H., H.S., K.Yagi, Y.O., T.K., M.d.H., P.C., J.W.S.; Funding acquisition: P.C.; Project administration: C.H., H.S., K.Yagi, S.K., Y.O., T.K., M.d.H., P.C., J.W.S.

## Declaration of interests

The authors declare that there is no conflict of interest.

## Methods

### RESOURCE AVAILABILITY

#### Raw data availability

The raw sequence files have been uploaded to the DNA Data Bank of Japan (DDBJ). They included 761 libraries of CAGE sequencing of iPS cells after specific knockdown of 123 lncRNAs and other control genes (DRA013844), RNA-sequencing with and without *EXOSC3* knockdown (DRA013847), Subcellular fractionation RNA-sequencing (DRA013845), RNA-sequencing of the untreated iPS cells (DRA013846), and RADICL-sequencing of the untreated iPS cells (DRA014412).

#### Processed data availability

https://fantom.gsc.riken.jp/6/suppl/Yip_et_al_2022/ contains supplementary tables and processed raw data.

Table S1: Results of knockdown and growth phenotype, Related to Figure 3

Table S2: Properties of 390 lncRNAs, Related to Figure 3

Table S3: ASO-pair-base and lncRNA-target-base molecular phenotype, Related to Figure 4

Table S4: Integration of Hi-C, RADICL-seq and eQTL to knockdown effects, Related to Figure 5

Table S5: LncRNAs in cell identities, Related to Figure 6

Table S6: LncRNA activities in HDF and iPS cells, Related to Figure 7

Table S7: Sequences of ASOs, siRNAs and primers, Related to STAR Methods

https://fantom.gsc.riken.jp/zenbu/reports/#FANTOM6_iPSC contains interactive webpage for visualizing the data.

## EXPERIMENTAL MODEL AND SUBJECT DETAILS

Human induced pluripotent stem (iPS) cells were generated by infecting human dermal fibroblast-fetal (HDF-f, Cell Applications INC.) with a non-integrating Sendai-based viral vector (SeV) coding for OCT3/4, SOX2, KLF4 and c-Myc, as described.^49^ The iPS cells were cultured in StemFit medium (ReproCELL Inc., Yokohama, Japan) under feeder-free conditions at 37°C in a 5% CO_2_ incubator. The iPS cells were authenticated by karyotyping, stem cell markers expression and teratoma formation were conformed in our previous study.^50^ The cells were plated on a culture dish pre-coated with laminin-511 (iMatrix-511: Nippi, Tokyo, Japan). Rho-associated kinase (ROCK) inhibitor, Y-27632 (Wako Pure Chemical Industries Ltd., Osaka, Japan) was added to the cells at a final concentration of 10 μM during the first 24 h of incubation after plating. StemFit medium is refreshed daily until the next passage. The cells were dissociated and detached by incubating with TrypLE select (Thermo Fisher Scientific) followed by scraping in StemFit medium. The cell viability in each passage is > 95%, as monitored by a trypan blue exclusion assay. For spontaneous differentiation conditions, FGF2 was removed from the complete medium 24 h after plating the cells.

Human dermal fibroblasts (HDFs) used were primary cells derived from the dermis of normal human neonatal foreskin cells (Lonza, catalog number: C2509). The cells were authenticated by Lonza. The cells were cultured in Dulbecco’s Modified Eagle’s medium (high glucose with L-glutamine) supplemented with 10% fetal bovine serum at 37°C in a 5% CO_2_ incubator. The passage number of the cells for transfection was maintained at six or seven.

### METHOD DETAILS

#### Gene models, lncRNA target selection and ASO design

The gene models and gene classes (divergent, antisense and intergenic lncRNAs) were interpreted and defined by FANTOM CAGE Associated Transcriptome (CAT).^1^ The selection of the 390 lncRNA targets cover different expression levels and different gene classes, while maintaining an expression level above 0.1 TPM according to the mean of three different iPS lines.^1^ Selection was achieved by manually curating the loci suitable for ASO design (e.g. without overlapping other mRNAs on the same strand).

The strategy of the locked nucleic acid (LNA) phosphorothioate gapmer ASO design has been described previously.^8^ Briefly, ASOs were designed with a central DNA gap flanked by 2-4 LNA nucleotides at the 5’ and 3’ ends of the ASOs. For each lncRNA target, we used the unspliced transcript sequence from FANTOM CAT as a template for designing a minimum of 5 ASOs per lncRNA. An exhaustive list of all target recognition sequences of 16-19 nt in length within the unspliced target lncRNA sequence was analyzed against the entire transcriptome to determine the prevalence of each *k*-mer, when allowing for 0 or 1 mismatched positions. Furthermore, the accessibility of each *k*-mer in the target sequence was estimated using RNAplfold using the ViennaRNA package.^51^ Subsequently, ASO designs were generated by applying various LNA/DNA design motifs to the reverse complement of the *k*-mer sequences. The LNA/DNA design patterns consisted of any combination of 2-4 LNA modifications at the 5’ and 3’ ends and central DNA window. For each ASO design, we calculated a series of characteristics, including predicted melting temperature for the ASO:RNA duplex, presence of problematic sequence motifs (G-quadruplex and other homopolymers, 5’ terminal GG) as well as propensity to fold/co-fold estimated by using RNAfold and RNAcofold (ViennaRNA package) on the ASO sequence. Finally, ASOs were selected based on properties of both the *k*-mer binding site in the target lncRNA sequence and characteristics of the different ASO designs, such that ASOs had zero perfectly matched off-targets, and no more than 1 imperfectly matched off-target, when allowing for 1 mismatched position. In addition, ASOs were selected to have a predicted melting temperature (T_m_) in the range of 50-56°C.^52^ A total of 2,225 ASOs targeting 390 lncRNAs were selected for the study. The ASOs were synthesized by Exiqon (1,923 ASOs) and GeneDesign Inc. (302 ASOs).

#### Transfection

The iPS cells were transfected in a 96-well plate. The cells were seeded 24 h before transfection at a density of 7,200 cells per well. A final concentration of 20 nM siRNA (Thermo Fisher Scientific) or ASO, and 0.2 µL lipofectamine RNAiMAX (Thermo Fisher Scientific) was mixed in 20 µL OptiMEM (Thermo Fisher Scientific). These transfection reagents were incubated at room temperature for 10 min and mixed with 90 µL StemFit medium without ROCK inhibitor. The medium from the 96-well plate was aspirated and replaced with the transfection mix. The volume and cell number were scaled in proportion if other sizes of well were used. The whole screening included 2,225 ASOs (390 lncRNA targets) in duplicate. Each 96-well plate included 2 wells for mock transfection (lipofectamine alone), 4 for non-targeting negative control (NC, Exiqon) and 2 for ASO targeting *MALAT-1* (MA-1, Exiqon) to control knockdown efficiency. For the transfection for HT-CAGE screening, 317 ASOs (123 lncRNA targets) were selected according to knockdown efficiency and growth phenotype in the screening. They were transfected with duplicates, in parallel with 4 untreated samples, 20 mock transfections, 48 NC and 8 MA-1. Several ASOs targeting characterized genes including *POU5F1*, *LncRNA-ES1*, *LncRNA-ES3*, and siRNAs targeting *POU5F1* and *EMC7* were also included as positive controls. The sequences of ASOs and siRNAs used in this study were listed in Table S7.

#### Real-time imaging

Phase-contrast images of transfected cells were captured every 3 h for 2 days with 4 fields per well by the IncuCyte live-cell imaging system (Essen Bioscience). The confluence in each field was analyzed by the IncuCyte software. The mean confluence of each well was taken from each time point starting from 15 h post-transfection until the mean confluence of the NC ASOs in the same plate reached 90%. The growth rate in each well was calculated as the slope of a linear regression. A normalized growth rate of each sample was calculated as the growth rate of the samples divided by the mean growth rate of the 4 NC ASOs from the same plate. Student’s *t*-test was performed between the growth rate of the duplicated samples and the 4 NC ASOs, assuming equal variance. Transfection of an ASO was considered to inhibit growth if the normalized growth rate < 0.63 (6 standard deviations from the mean normalized growth rate of NC) and *t*-test *p*-value < 0.05 (comparison between sample and NC).

#### RNA purification

The harvested lysates were subjected to purification using the RNeasy 96 Kit (Qiagen) and epMotion automated liquid handling systems (Eppendorf) according to the manufacturers’ instructions. The eluted RNA was quantified by the NanoDrop™ spectrophotometry platform (Thermo Fisher Scientific). The purified RNA samples were stored at −80 °C until they were used for the RT-qPCR assay or the HT-CAGE library construction.

#### Real-time quantitative RT-PCR

Real-time quantitative RT-PCR was performed by One Step PrimeScript™ RT-PCR Kit (Takara Bio Inc.) using the epMotion automated liquid handling system (Eppendorf). For each lncRNA target, 3 primer pairs against the specific lncRNA target were used. The expression level was normalized by *GAPDH* while the knockdown efficiency was calculated from the fold-change between specific samples and the transfection of NC ASOs. The knockdown efficiency of *MALAT1* was monitored along each run and across all the runs where all the samples have >90% KD. The primers’ sequences are listed in Table S7.

#### WST-1 growth assay

Cells were transfected in a 96-well format as described above. At 48 h post-transfection, the medium was discarded and replaced with 100 µL StemFit medium and 10 µL WST-1 solution (Abcam). The cells were incubated at 37°C in a 5% CO_2_ incubator for 1 h, followed by measurement of OD 450 in a microplate reader. All absorbance values were subtracted by the control value from empty wells containing 100 µL StemFit medium and 10 µL WST-1 solution. These subtracted absorbance values were further normalized by cells transfected with NC ASOs seeded on the same plate. This batch of transfection included 317 ASOs selected for HT-CAGE. These transfected cells have real-time imaging measurement in parallel.

#### Colony formation assay

Reverse transfection was used for this assay. The cells were incubated with ROCK inhibitor in complete StemFit medium for 1 h before detachment by Accutase. After detachment, cells were centrifuged and resuspended in StemFit medium. Transfection mix was prepared as a 200 µL solution containing 2 µL lipofectamine RNAiMAX and 20 nM siRNA / ASO in the StemFit medium. The mix was maintained at room temperature for 10 min, followed by addition of 10,000 cells in 200 µL StemFit medium. After 15 min, 3,000 cells in 0.5 mL StemFit medium were seeded onto a 24-well pre-coated with laminin-511. Cells were refreshed after 3 days and stained by 0.5% crystal violet at day 6. The wells were imaged by the Celigo Image Cytometer (Nexcelom Bioscience) and the number of colonies were counted by ImageJ.^53^

#### High-throughput low quantity cap analysis of gene expression (HT-CAGE)

We selected 317 ASO knockdown samples from 123 lncRNA targets for HT-CAGE analysis. The selection was done by manual curation where we selected ASOs with relatively similar knockdown efficiency, where 301 are effective ASOs and 16 of them show marginal knockdown efficiency (> 40% in general). Among the 123 lncRNAs, 20 show growth phenotype from the cellular phenotyping. In parallel, NC ASOs, negative non-targeting control and positive controls such as ASO and siRNA targeting *POU5F1* were also subjected for HT-CAGE.

Fifty nanogram of purified RNA was used from each sample, which was 96-multiplexed for constructing one single-strand CAGE library,^22^ according to an improved version of nAnT-iCAGE protocol^21^ that avoids any bias from PCR amplification. Libraries were subjected to 50-base pair-end sequencing using an Illumina HiSeq 2500 instrument. Tags were de-multiplexed and mapped to the human genome (hg38) using STAR.^54^ The median number of mapped CAGE transcription start sites (CTSS) obtained from each sample is about 500,000. Gene expression profile was analyzed by intersecting^55, 56^ the mapped CTSS with the permissive CAGE peaks regions (of 709,716 promoters) annotated in FANTOM CAT.^1^ Promoter-level expression profiles were used in motif activity response analysis (MARA). Gene-level expression profiles were aggregated from promoter-level profiles for other downstream analyses.

#### Global analysis of cellular phenotype of same-target ASOs

From 2,225 ASOs, 750 ASOs showed effective knockdown (knockdown efficiency > 50% in one of the 3 primer sets) while 1,475 of them were considered without effective knockdown. The one-way ANOVA test was carried out on the 200 lncRNAs targets from 686 ASOs, with 2-10 successful ASOs per target, using the normalized growth rate for each ASO and gave the *p*-value = 2.79 × 10^-5^ (**Fig. 3E**). The analysis was done in R 4.0^57^ using the ‘aov’ function (‘*stats*’ 4.1.1 package) and the target labels were permuted using the ‘shuffle’ function (‘*mosaic*’ 1.8.3 package). Among the 200 lncRNAs with ≥ 2 effective ASOs (successful KD), we paired the 750 ASOs to those targeting the same lncRNA and resulted in 1,036 ASO-pairs. We used the 1,475 ASOs without effective knockdown as the background. The growth inhibition effects mediated by these ASOs are the maximum possible background effects caused by side effects (toxicity or off-target effect) of ASO-transfection. These ASOs were randomly paired for 1,036 ASO-pairs for 10,000 times. By considering concordant growth inhibition (growth inhibition from both ASOs in a pair) as a hit, 6.4% of the same-target-ASO-pairs resulted as hits, that is higher than the background pairs in all 10,000 randomizations (empirical p-value < 0.0001, **Fig. S1C**).

#### Growth phenotype determination of lncRNA target

At a robust level, lncRNAs with ≥ 2 independent effective ASOs resulted as growth inhibition (normalized growth rate < 0.63, t.test *p*-value < 0.05) are considered as showing growth phenotype (36/200). To implement a stringent level with statistical correction, we applied binomial and required *p*-value < 0.05 as a stringent cutoff. From the 2,225 ASOs, 1,475 of them were considered without effective knockdown. We set these 1,475 ASOs as the background to represent the maximum possible chance of growth inhibition mediated by side effects. The rate of significant growth inhibition from the background and for the effective ASOs is 15.2% and 23.4%, respectively. Since the number of successful knockdown ASOs for each lncRNA is different, we set up a binomial test to assess how many ASOs showing growth inhibition is significantly higher than background (binomial *p*-value < 0.05). Briefly, we have taken 15.2% as background growth inhibition, with the chance of ≥ 2 ASOs in the group of lncRNA with 2 ASOs showing successful knockdown is 0.152^2^ = 0.023104, which means that a false positive occurring randomly is < 5%. We set a conditional cutoff for different groups of lncRNA having different numbers of successful knockdown ASO to keep the chance of false positives lower than 5%. Using this stringent cutoff, 11 lncRNAs were considered to have a growth phenotype.

#### Correlation between gene properties and growth phenotype

These analyses were done target-based. LncRNAs were grouped as successful knockdowns if ≥ 2 ASOs showed reduction of the specific RNA level by 50%, considering the primer set showing the strongest knockdown result from the RT-qPCR. By this criterion, 200 lncRNAs were grouped as successful knockdown and 190 were grouped as no KD. From the 200 lncRNAs, 36 of them were grouped as growth phenotypes where ≥ 2 ASOs showed reduced normalized growth rate compared to that of the NC ASOs significantly with a cutoff of smaller than 0.63 (6 standard deviations from the mean of NC). The 165 lncRNAs were grouped as having no growth phenotype. The tests were performed between successful knockdown (n=200) and no knockdown (n=190), and between growth phenotype (n=36) and no growth phenotype (n=164). Discrete gene properties, including derived from promoter or enhancer, intergenic or non-intergenic, antisense or non-antisense, divergent or non-divergent, were accessed by Fisher’s exact one-tailed test. Gene properties with continuous values were analyzed by the Wilcoxon one-tailed test. The transcription initiation regions (TIR) and exon conservation scores were derived by rejected substitution scores as described.^1^ Exosome sensitivity score was derived from comparing the transcriptome of control and EXOSC3 knockdown in our iPS line using the formula (TPM_Exosome-suppressed_ - TPM_Control_) / TPM_Exosome-suppressed_.^58^ Expression levels of the targets were revealed from CAGE sequencing and RNA-seq data of our iPS line. The number of exons refers to the maximum number of exons from any transcripts of the lncRNAs. Cell type specificity (Gini index calculated from 71 cell types in FANTOM5) is described in the corresponding methods. The number of Hi-C contacted, eQTL-contacted and RADICL contacted protein coding genes with TPM > 0.1 are identified as described in the corresponding methods. Comparison based on growth phenotype with stringent cutoff (11 VS 189) and molecular phenotype (36 VS 87) are shown in Table S2. The Wilcoxon tests were done in R 4.0 using the wilcox.test function (‘*stats*’ 4.1.1 package).

#### Motif Activity Response Analysis (MARA)

MARA^59^ was performed using TMM normalized and batch corrected promoter expression profiles for all the ASO knockdown and controls (NC ASOs and other positive controls) libraries (732 CAGE libraries). Batch correction was performed by the “removeBatchEffect” function in the limma R package. All promoters with expression ≥ 1 TPM at least in 70% CAGE libraries (13,883 promoters) were used for the analysis. Transcription factor binding sites (TFBS) for hg38 were predicted as described previously^60^ using MotEvo^61^ for the set of 190 position-weight matrix motifs in SwissRegulon (released on 13 July 2015)^62^ on a multiple alignment of genome assemblies. The number of predicted TFBS were counted for each motif in the −300 to +100 base pair from the midpoint of the FANTOM CAT promoters. Next, MARA was performed to decompose CAGE expression profiles of the promoters in terms of their associated motifs, yielding the activity profile of all the motifs with at least 150 TFBS associated with the expressed promoters across the iPS cells knockdown and control samples. The comparison was done by 2 knockdown libraries per ASO versus 47 libraries of NC ASOs by using the Wilcoxon test for the *p*-value, which was adjusted by the Benjamini-Hochberg method. Net activity was calculated by the mean activity of each ASO minus the mean activity of the 47 NC ASOs. The motifs showing FDR < 0.1 were considered significant.

#### Differential expression

Differential expression of 10,868 genes was performed by *edgeR* (ver. 3.10)^63^ after aggregating the promoter-level profiles. A design of 2 libraries of ASO versus 47 libraries of NC using TMM normalization was adopted after batch effect correction. FDR was obtained by p.adjust from the p-value of *edgeR* using the Benjamini-Hochberg method. Across sample Z-score was calculated from the log_2_FC of a gene across the 317 ASO knockdown samples. The genes showing FDR < 0.1 and |Z-score| > 1.645 were considered as significant changes for downstream analyses. The raw data are deposited (DDBJ: DRA013844) while the tables for raw count and the TMM-normalized count per gene (n = 10,868) per library (n = 732) are available in here: https://fantom.gsc.riken.jp/6/suppl/Yip_et_al_2022/.

#### Gene set enrichment analysis (GSEA) for cellular pathways

Gene set enrichment (GO Biological Process, GO Molecular Function, GO Cellular Component, KEGG, Hallmark and Reactome) analysis was carried separately for each set of pathways and each ASO knockdown using ‘*fgsea*’ (1.8.0) R-package^64^ with the following parameters: minSize=15, maxSize=1000, nperm=10000, nproc=1. Each pathway was tested using gene ranks based on -log_10_(*p*-value) × sign(log_2_FC) from the *edgeR* analysis. Across samples, the Z-score was obtained by NES dividing into positive values and negative values from the 317 ASO samples. Pathway was considered significant if adjusted *p*-value < 0.05 and |Z-score| > 1.645.

#### Correlation between GSEA pathways and cellular phenotype

From the *fgsea* result, we extracted the -log_10_(adjusted *p*-value) × sign(NES) of the 317 ASOs for each pathway. These values were compared with the normalized growth rate of the 317 ASOs by Spearman correlation for each pathway. The rho values for all the pathways are reported in **Figure 4B**.

#### Degree of transcriptome change predicts growth phenotype

The 317 ASO knockdown profiles from *edgeR* were compared with the NC ASO-background by their -log_10_(*p*-value) of all the 10,868 genes. Briefly, a KS curve was plotted for each knockdown profile to calculate the mean distance between one ASO and the NC-background. Each NC-ASO-combination was also compared with the NC-background. Next, ASOs were paired if targeting the same lncRNA. The calculated distances of the two ASOs were compared with distances of 47 NC ASOs by the Wilcoxon test. The *p*-values from the tests were adjusted by the Benjamini-Hochberg method across all the ASO-pairs for FDR. The mean distance of the 2 ASOs and the -log_10_(Wilcoxon *p*-value) were plotted in the chart (**Fig. 4A**). The source data can be found in Table S3.

#### Reciprocal enrichment test for ASO-pairs

We performed a reciprocal enrichment test to identify concordant ASO-pairs, which implies a consistent change of molecular phenotype between two ASOs targeting the same lncRNA. The method of the enrichment is illustrated in **Fig. S3A-C**. Briefly, we rank the knockdown profile for each ASO using -log_10_(p-value) × sign(log_2_FC) from *edgeR*. The top and bottom 25, 50, 75, 100, 125, 150, 175 and 200 genes were taken as genesets. These genesets for all the 317 ASOs were tested for enrichment against all the knockdown profiles of the 317 ASOs by ‘*fgsea*’ (1.8.0) where each geneset size was tested separately for the adjusted *p*-values. The NES values of the different-target-ASO-pairs were scaled for a normalized background distribution. A same-target-ASO-pair was considered significant if both the forward test and the reverse test adjusted *p*-values < 0.05 with |NES Z-score| > 1.645. Both positive and negative enrichment was considered a hit. The hits were called from the union of any of the geneset sizes (**Fig. 4C**).

#### Concordance test for differentially active features

The 317 ASOs were paired for targeting the same lncRNAs and resulted in 293 same-target-ASO-pairs, which were then shuffled and enforced into different-target-ASO-pairs, 10,000 times for permutation. Here, we use the *edgeR* results as an example. To determine if ASO knockdown resulted in a specific effect stronger than noise (effects of ASO transfection), we used the Jaccard index of common significant DEG in the same direction to quantify the degree of concordance between two ASOs in a pair. Briefly, differentially expressed genes (excluded the target lncRNAs) from each ASO-knockdown were taken with FDR < 0.1 and |Z-score| > 1.645. From the 293 ASO-pairs, the mean Jaccard index was calculated for the same-target group. The mean values for each permutation of the different-target group were calculated to generate a distribution. The empirical *p*-value represents the chance that the mean values of the same target group is not greater than the different-target group. For all the tested conditions using the Jaccard index in *edgeR*, the empirical *p*-values are < 0.0001.

#### Optimal cutoffs of molecular features

To improve the performance, we attempted to apply two types of background correction, NC-background correction and across sample (317 ASOs) background correction. For the NC-background, maximum combinations of two libraries from 47 NC ASOs CAGE libraries were obtained. These 1,081 NC combinations were applied to *edgeR*, *fgsea* and MARA analysis similarly as the ASO samples. The resulting log_2_FC, NES and motif activity values of these 1,081 combinations were scaled as normalized backgrounds. For the across sample background, log_2_FC, NES and motif activity from the 317 ASO samples were scaled as normalized background. The transformed Z-score of the actual values were used as background correction using |Z-score| > 1.645 as a cutoff.

After the assessments by using Jaccard indices (**Fig. S3F**), all the tested conditions in all the molecular features revealed the same-target-ASO-pairs are more concordant than the different-target background significantly (empirical *p*-values are < 0.0001). We selected the best cutoffs by maximizing the ratio of concordance of same-target-ASO-pairs to different-target-ASO-pairs, resulted in FDR < 0.1 with |across-sample Z-score| > 1.645 for genes (*edgeR* analysis), adjusted *p*-value < 0.05 with |across-sample Z-score| > 1.645 for pathways (*fgsea* analysis) and FDR < 0.1 for motifs (MARA analysis).

After determining the cutoffs for the differentially active features (DEG, pathways & motifs), the numbers of common features between two ASOs in the ASO-pairs were collected. From the concordant ASO-pairs, we require at least one common feature and resulted in the final 36 concordant lncRNAs.

#### Isoform coverage similarity for ASO-pairs

In this analysis, only the final 36 concordant lncRNA targets were included. From these 36 concordant lncRNA targets, there are 102 ASOs involved, resulting in 111 same-target-ASO-pairs. Among them, 59 pairs are significantly concordant. To identify isoform targets, we mapped the ASOs to all transcripts defined by FANTOM CAT annotation^1^ and excluded those transcripts that were absent in the RNA-seq data of iPS cell line. The Jaccard indexes considering the number of overlap and distinct transcripts targeted by the 2 ASOs in a pair were calculated. We then compared the Jaccard indexes between the “not concordant group” and the “concordant group” by using one-tailed Wilcoxon Test. Further, from our RNA-seq data we extracted the TPM for each transcript variant and weighted each transcript before calculating the Jaccard index. These weighted Jaccard indexes from the “not concordant group” and the “concordant group” were compared by using one-tailed Wilcoxon Test.

#### Hi-C

The experimental setup of Hi-C for iPS cells is the same as HDF cells, as described previously.^8^ The analytical pipeline has been described.^26^ Briefly, after the binomial test at 10 kb resolution for intra-chromosomal interactions, respectively, interacting regions with FDR < 0.05 were considered significant. The bin regions were intersected with the whole gene regions (gene region + 800 bp upstream the transcription start site) of the lncRNA targets, the whole gene regions of mRNAs and the promoter regions of mRNAs (800 bp upstream and 200 bp downstream of the transcription start site of all transcripts) defined in FANTOM CAT.^1^ The whole gene region against whole gene region interaction was used to identify contacted protein coding genes of the lncRNA targets. We used the contacted protein coding genes that are detectable in HT-CAGE to construct genesets for enrichment analysis against the knockdown profiles of each ASO by ‘*fgsea*’ (1.8.0). The whole gene region of lncRNA against the promoter region of mRNA interaction was used as an extra indicator of regulation. The TAD regions in the whole genome of iPS cells were predicted by the Hi-C data at 50 kb resolution, as described previously.^26^ Inter-chromosomal Hi-C interactions was identified at 50 kb resolution and mapped to genes the same way as intra-chromosomal interactions. The inter-chromosomal Hi-C interaction was only used to filter RADICL inter-chromosomal signal. However, there were only 62 lncRNA-protein coding genes inter-chromosomal Hi-C pairs identified and they did not overlap with any RADICL signals.

#### Expression quantitative trait loci (eQTLs)

The eQTL-linked genes were obtained from the Genotype-Tissue Expression (GTEx) project (v8).^27^ The eGene and significant variant-gene associations based on permutations were taken from 49 tissues with the permutation *q*-value < 0.25. The genomic location of the genetic variants was intersected with the 390 lncRNA targets for their whole gene region (gene region + 800 bp upstream the transcription start site). For the eGenes, only protein coding genes having expression with at least 0.1 TPM from iPS cells or being detected by HT-CAGE were included. From the 36 concordant lncRNAs, 31 of them showed at least one significant eGene.

#### RADICL-seq

RNA And DNA Interacting Complexes Ligated and sequenced (RADICL-seq) was performed, as described previously.^24^ The analytical pipeline was performed as described previously^26^ with modification. Briefly, RNA mapped to a defined set of small RNAs was removed from the dataset. The genome was then divided into 25 kb bins and the annotated RNA reads were aggregated. The background probability for a bin was calculated by dividing the count of trans mRNA binding in that bin by the total number of trans mRNA reads. To estimate the significance of binding each gene in every bin, we performed a one-sided binomial test using binom_test (x,n,p) from scipy where x = the number of reads for the gene in the bin; n = total number of reads for the gene; p = background probability calculated using trans-binding mRNA in the bin. FDR was obtained by p.adjust from the binomial *p*-values using the Benjamini-Hochberg method across all the bins per RNA. The regions with at least 3 reads and FDR < 0.05 were considered significant. The regions from the DNA side were intersected towards the whole gene regions (gene region + 800 bp upstream the starting site) of mRNAs and the promoter regions of mRNAs (800 bp upstream and 200 bp downstream of the CTSS of all transcripts) defined in FANTOM CAT.^1^ The RADICL-contacted protein coding genes for each of the lncRNA target were taken if the mRNA expressed higher than 0.1 TPM in iPS cells or being detected in HT-CAGE. RADICL-contacted genes with the minimum genomic distance greater than 1 Mb and without significant Hi-C interaction (intra- and inter-chromosomal) were defined as *trans*-interaction. The identification of significant DNA binding wad performed in all 390 lncRNA targets. Within the 36 concordant lncRNAs, 35 showed significant *cis*-interaction and 17 showed *trans*-interaction with at least 1 protein coding gene.

#### Global enrichment of contact and significant DEGs

All the lncRNAs were paired with 9974 protein coding genes that are detectable in HT-CAGE in iPS cells. For all these pairs we extracted the *edgeR* results of the most significant ASO to define significant DEGs, and compare this to the presence of *cis*-interactions and trans-interactions. We only used the pairs derived from the 35 lncRNAs showing at least one *cis*-interaction, the 17 lncRNAs showing at least one *trans*-interaction (no significant Hi-C contact and at least 1 Mb genomic distance) and the 31 lncRNAs showing at least one eGene for the respective tests. Fisher’s exact one-tailed test was performed.

#### RNA-DNA sequence complementary analysis

The transcript models with the highest expression in the soluble nuclear fraction were used to represent the sequence of the lncRNA targets. These RNA sequences were mapped with the whole gene regions and the promoter regions of the RADICL-contacted mRNA using a window size of 20 bp allowing 20% mismatch. The mapping was done by the TDF software.^65^ Any interaction with the whole gene regions (gene region + 800 bp upstream the starting site) of mRNA located > 1 Mb from the lncRNA gene region was counted.

#### Enrichment analyses between knockdown profile and contacts

The enrichment tests of protein coding genes showing Hi-C contacts, RADICL-seq contacts and eQTL linkages with any of the 36 concordant lncRNAs were performed with ‘*fgsea*’ (1.8.0) R-package.^64^ The 9974 protein coding genes were ranked according to the -log_10_(*p*-value) × sign(log_2_FC) from the *EdgeR* analysis of each ASO. Enrichment was tested for the connected coding genes in Hi-C, RADICL-seq or eQTL towards the ranked profile. The *p*-values were then the adjusted by the Benjamini-Hochberg method. Inclusion criteria of protein coding genes for Hi-C contacts: *q*-value < 0.05; RADICL-seq contacts: *q*-value < 0.05, read>=3; eQTL linkages: *q*-value < 0.25.

#### Cell type specificity on expression level

FANTOM 5 CAGE data on ∼120K genes with 384 samples with cell ontology classification^58^ plus iPS and ES cells (iPS in this study was not included) were used to determine cell type specificity for all the RNA including the lncRNA targets. CAGE libraries used were listed in Table S5. We applied the Gini index for each gene from the RLE normalized TPM across the 71 cell types. To compare cell type specificity between mRNA and lncRNA, genes were grouped by the maximum TPM across the 71 cell types. Gene annotation of mRNA and lncRNA was according to the FANTOM CAT gene classes.^1^ Statistical significance in each group was acquired by applying the Student’s *t*-test where *p*-value < 0.05 was considered significant in **Figure 7A**.

For building a geneset for the 71 cell types for cell type enrichment analysis, expression levels of a gene in a cell type lower than 5% of the maximum TPM were considered as lack of expression, and the corresponding Gini index set as 0. Only including the 10,868 genes detected from HT-CAGE, the top 25, 50, 75, 100, 125, 150, 175 and 200 genes ranked by the Gini index were taken for each cell type. These genesets for 71 cell types were tested for enrichment against all the knockdown profiles of the 317 ASOs by ‘*fgsea*’ (1.8.0) where each geneset size was tested separately for the adjusted *p*-values. The NES values across the 317 ASOs were scaled and cross-samples Z-scores were obtained. LncRNA targets having at least 2 ASOs with adjusted *p*-value < 0.05 and |Z-score| > 1.645 towards a cell type were considered significant. A list of significant lncRNA targets was obtained from the union of the 8 geneset sizes.

#### Cell fractionation for RNA-sequencing

Subcellular fractionation of native iPS cells was performed similarly as HDF as previously described^8^ with minor optimization according to the cell type. Briefly, approximately 10 million cells were used per fractionation experiment to obtain cytoplasmic (CYTO), soluble nucleoplasmic (SNUC) and chromatin (CHR) fractions. Trypsinized cells were washed and lysed using a cold lysis buffer containing 0.15% Igepal CA-630, 10 mM Tris pH 7.5, 150 mM NaCl. The lysate was centrifuged in a sucrose cushion, after which the supernatant was taken as the CYTO fraction. The nuclear pellet was washed once in a buffer containing 20 mM HEPES pH 7.5, 50% glycerol, 75 mM NaCl, 1 mM DTT, 0.5 mM EDTA and resuspended in the same buffer. An equal volume of nuclear lysis buffer containing 20 mM HEPES pH 7.5, 300 mM NaCl, 1M Urea, 1% Igepal CA-630, 10 mM MgCl_2_, 1 mM DTT, 0.2 mM EDTA was added and incubated on ice for 5 min. After centrifugation, the supernatant was considered as the SNUC fraction and the pellet as the CHR fraction. The chromatin pellet was washed once in a buffer containing 10 mM HEPES pH 7.5, 10 mM KCl, 10% glycerol, 340 mM sucrose, 4 mM MgCl_2_, 1 mM DTT and resuspended in the same buffer. RNA from each fraction was isolated using Trizol LS (Invitrogen), according to the manufacturer’s instructions. Next, DNase I treatment followed by phenol-chloroform extraction was conducted. The volume and concentration of the RNA isolated from each fraction was recorded, with an equal amount of RNA for each fraction was used for RNA-sequencing.

Single-end read RNA-sequencing was performed with the Illumina platform by HiSeq2000. Fastq data were mapped towards human genome assembly GRCh38 (hg38) and FANTOM CAT permissive transcript model^1^ using TopHat2.^66^ The resulting gene length and library size normalized tpm values in each fraction were multiplied by the corresponding RNA content factors (amount of RNA extracted from each fraction divided by the sum of the RNA amount of all fractions). The fraction-normalized tpm (nTPM) was used to calculate the nucleus ratio from the whole cell ((SNUCnTPM + CHRnTPM) / (CYTOnTPM + SNUCnTPM + CHRnTPM)) and the chromatin ratio from the whole nucleus (CHRnTPM / (SNUCnTPM + CHRnTPM)).

The median value of the nucleus / whole cell ratio was calculated from all the genes with >1 TPM in iPSC (as 0.1418). Targets with the nucleus ratio smaller than this were considered as cytoplasmic-enriched. The remaining targets were grouped as chromatin-enriched for their chromatin / whole nucleus ratio greater than the median. The remaining targets are soluble nucleoplasmic enriched.

#### Concordant knockdown responses between HDF and iPS cells

The knockdown profiles by individual ASO in iPS cells were generated by *edgeR* as earlier described. The knockdown profiles in HDF cells were obtained by DESeq2^67^ after nAnTi-CAGE sequencing, as described previously.^8^ Only genes (n=10,054) overlapping in both analyses were included, with these genes ranked by -log_10_(*p*-value) × sign(log_2_FC). Genesets with sizes of 25, 50, 75, 100, 125, 150, 175 and 200 were generated from each ASO knockdown profile from the top and the bottom of the ranking. Gene sets were tested for enrichment by ‘*fgsea*’ (1.8.0) in the knockdown profiles of the ASO with the same targets in the other cell type. Enrichment tests were also performed towards the knockdown profiles from the different-target-ASO-pairs (n = 250, 10,000 permutation) to build the background. In total, 250 different-cell-types-same-target-ASO-pairs were tested and significant pairs meet the following criteria in either up-/down-regulation in any geneset size: both forward and reverse enrichment adjusted *p*-values < 0.05 while both forward and reverse |different-target different-cell ASO-pairs background Z-score| > 1.645.

For the common significant molecular features applied to each same-target-ASO-pair, we used the same cutoffs defined above for iPS cells. For HDF, we used FDR<0.01, |log_2_FC| > 0.5 and |Z-score|>1.645 for genes; adjusted *p*-value < 0.05, |NES|>1 and |Z-score|>1.645 for pathways; FDR<0.05 and |Z-score|>1.645 for motifs. The target lncRNAs were excluded from the common differentially expressed genes.

## QUANTIFICATION AND STATISTICAL ANALYSIS

The real time imaging included 2 biological replicates for lncRNA ASO knockdown and 4 biological replicates for NC ASO knockdown. The mean confluence of the 4 imaging fields was taken for each replicate. Student’s *t*-test was performed between the growth rate of the duplicated samples and the 4 NC ASOs, assuming equal variance. Transfection of an ASO was considered to inhibit growth if the normalized growth rate < 0.63 (6 standard deviations from the mean normalized growth rate of NC) and *t*-test *p*-value < 0.05 (comparison between sample and NC). At a robust level, lncRNAs with ≥ 2 independent effective ASOs resulted as growth inhibition are considered as showing growth phenotype (36/200). At a stringent level, we applied a binomial test and required number of growth inhibition ASOs greater than the binomial cutoff at *p*-value < 0.05. Colony formation assay was performed with 3 biological replicates. Student’s *t*-test was performed between the lncRNA ASO knockdown and the NC ASO, assuming equal variance.

For the global analysis of cellular phenotype from the same-target ASOs, one-way ANOVA test was carried out on the 200 lncRNAs targets from 679 ASOs, with 2-10 successful ASOs per target, using the normalized growth rate for each ASO.

Enrichment of lncRNA target properties was performed by either Fisher’s Exact or Wilcoxon depends on the target properties shown as continuous or discrete values. All tests are performed as one-tail for the *p*-values.

For MARA, the comparison was done by 2 knockdown libraries per ASO versus 47 libraries of NC ASOs by using the Wilcoxon test for the *p*-value, which was adjusted by the Benjamini-Hochberg method. The motifs showing FDR < 0.1 were considered significant.

For differential expression, FDR was obtained by using the Benjamini-Hochberg method from the *p*-value calculated by *edgeR*. Across sample Z-score was calculated from the log_2_FC of each gene across the 317 ASO knockdown samples. The genes showing FDR < 0.1 and |Z-score| > 1.645 were considered as significant changes

For gene set enrichment assay, each pathway was tested using genes ranked by -log_10_(*p*-value) × sign(log_2_FC) derived from the *edgeR*. Analysis was performed separately for each set of pathways and each ASO knockdown using ‘*fgsea*’ (1.8.0)^64^ with the following parameters: minSize=15, maxSize=1000, nperm=10000, nproc=1. The Z-score was obtained by NES dividing into positive values and negative values from the 317 ASO samples. Pathway was considered significant if adjusted *p*-value < 0.05 and |Z-score| > 1.645.

For the degree of transcriptome change, ASO knockdown profiles from *edgeR* were compared with the NC ASO-background by their -log_10_(*p*-value) of all the 10,868 genes. The ASOs were paired if targeting the same lncRNA. The calculated distances of the two ASOs were compared with distances of 47 NC ASOs by the Wilcoxon test. The *p*-values from the tests were adjusted by the Benjamini-Hochberg method across all the ASO-pairs for FDR. Same-target-ASO-pairs with FDR < 0.05 and distance > 99.9% were considered as significantly concordant.

For the reciprocal enrichment test, we rank the knockdown profile for each ASO using -log_10_(p-value) × sign(log_2_FC) derived from *edgeR*. The top and bottom 25, 50, 75, 100, 125, 150, 175 and 200 genes were taken as genesets. These genesets for all the 317 ASOs were tested for enrichment against all the knockdown profiles of the 317 ASOs by ‘*fgsea*’ (1.8.0) where each geneset size was tested separately for the adjusted *p*-values. The NES values of the different-target-ASO-pairs were scaled for a normalized background distribution. A same-target-ASO-pair was considered significant if both the forward test and the reverse test adjusted *p*-values < 0.05 with |NES Z-score| > 1.645. Either positive or negative enrichment was considered as a hit.

For the cell type enrichment analysis, the top 25, 50, 75, 100, 125, 150, 175 and 200 genes ranked by the Gini index were taken for each cell type. These genesets for 71 cell types were tested for enrichment against all the knockdown profiles of the 317 ASOs by ‘*fgsea*’ (1.8.0) where each geneset size was tested separately for the adjusted *p*-values. The NES values across the 317 ASOs were scaled and cross-samples Z-scores were obtained. LncRNA targets having at least 2 ASOs with adjusted *p*-value < 0.05 and |Z-score| > 1.645 towards a cell type were considered significant. A list of significant lncRNA targets was obtained from the union of the 8 geneset sizes.

Concordant knockdown response between HDF and iPS cells was determined by reciprocal enrichment analysis. Significant concordant pair was defined as both forward and reverse enrichment adjusted *p*-values < 0.05 while both forward and reverse |different-target different-cell ASO-pairs background Z-score| > 1.645.

## SUPPLEMENTARY TABLES AND RESOURCES

https://fantom.gsc.riken.jp/6/suppl/Yip_et_al_2022/

---

**Figure S1.**
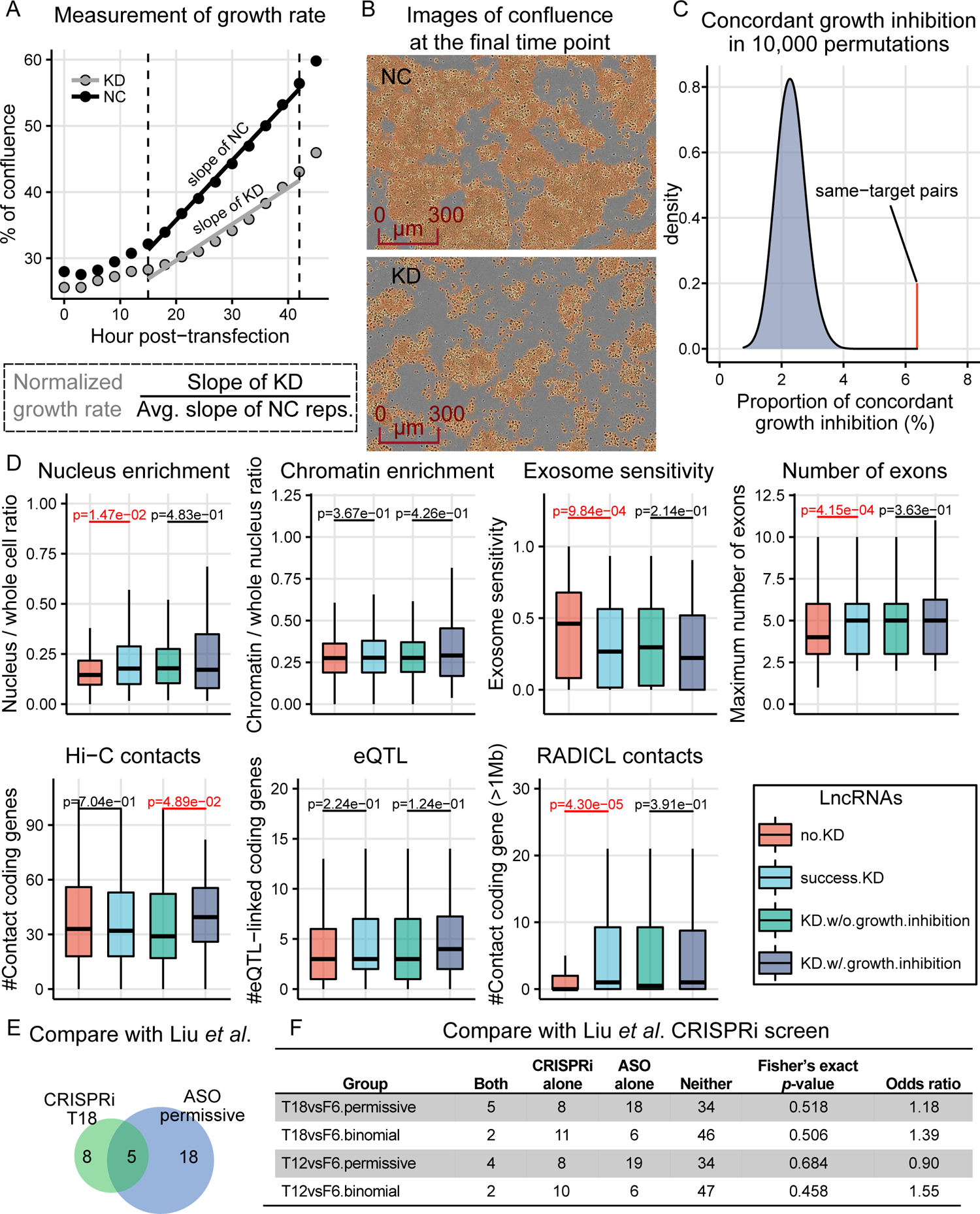
Cellular phenotypes and their correlations with genomic properties of lncRNAs, Related to Figure 3. **A,** Growth rate of each KD sample was normalized by NC ASOs from the same set of experiment. **B,** Example images of NC and KD taken at the final time point are shown. **C,** The empirical *p*-value for the chance that the number of concordant ASO-pairs is higher among the same-target ASO-pair than that of the different-target permutation is <0.0001. The background was generated by randomly pairing the “no KD” ASOs into 1036 ASO-pairs for 10,000 times. **D**, LncRNAs with successful KD are more enriched in the nucleus, less exosome sensitive and have a number of exons. LlncRNAs with a growth phenotype show greater number of exons and greater number of Hi-C contacts (lncRNA targets, n=390; noKD, n=190; KD, n=200; KD.growth, n=36; KD.no.growth, n=164). *p*-value < 0.05, highlighted in red. **E**, Venn diagram showing the growth related lncRNAs from the 65 common lncRNAs targeted in the CRISPRi screen by Liu *et al*. (1) and from this study. **F**, Fisher’s exact tests compare the growth phenotype between the Liu *et al*. (1) CRISPRi screen (2 time points) and this study (2 cutoffs).

**Figure S2.**
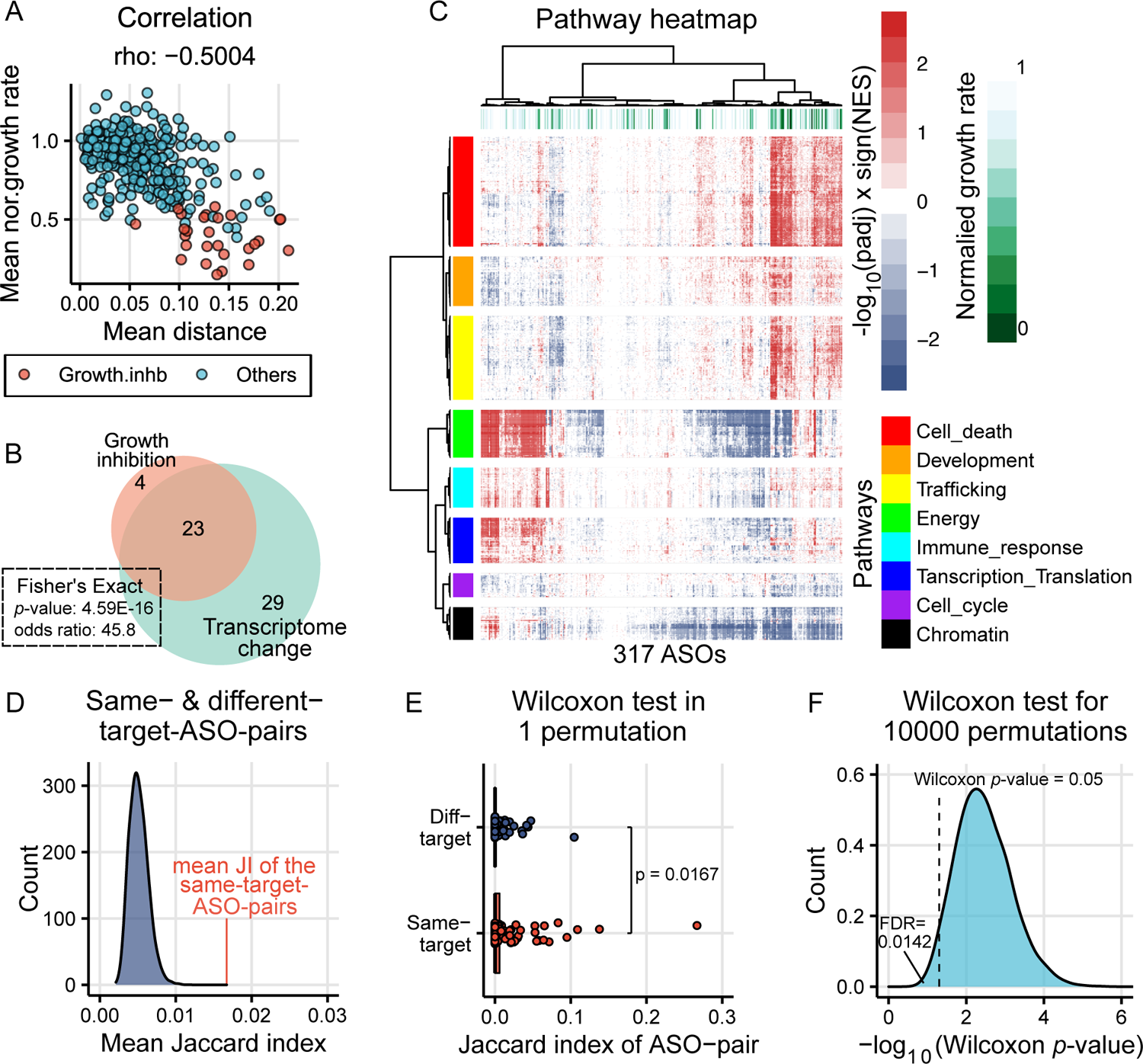
Correlation between molecular phenotype and cellular phenotype and lncRNA target specific results, Related to Figure 4. **A**, Correlation between global change and normalized growth rate for the 293 ASO-pairs. **B**, Venn diagram showing the significant ASO-pair hits of transcriptome change and growth inhibition. **C**, Heatmap from pathway *fgsea* analysis showing 317 ASOs. **D**, The distribution of mean Jaccard indices from the different-target-ASO-pair background and the mean Jaccard index of the same-target-ASO-pairs. Using the *edgeR* dataset using the cutoff of FDR < 0.1 & |across sample Z-Score| > 1.645, DEGs passing the cutoff from each ASO-knockdown were taken for 293 ASO-pairs while the same ASOs were permuted 10,000 times enforcing different-target pairing. The mean Jaccard indices were calculated for the same-target group and for each permutation of the different-target group. The empirical p-value (<0.0001) represents the chance that the mean of the Jaccard index of the same target group is not greater than the different-target group. **E**, Using the Jaccard index of overlap DEGs, non-redundant ASO-pairs (n=137) were selected randomly and the selected ASOs were permuted into different target pair for Wilcoxon test. The chart shows the result of one of the permutations. **F**, The p-value was recorded in each permutation and illustrated as -log_10_ *p*-value. False discovery rate of 0.0142 was calculated as 142/10,000 times where

**Figure S3.**
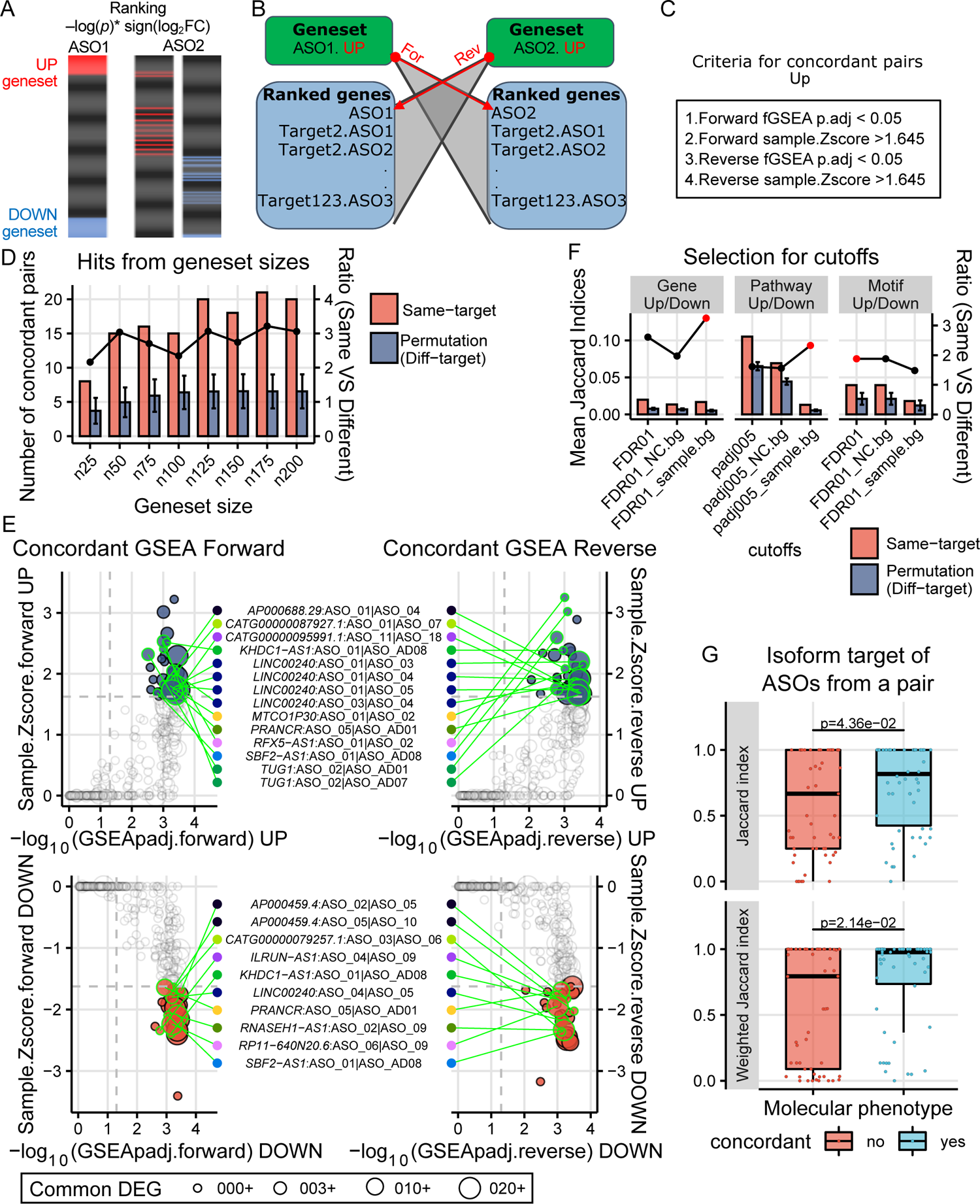
Knockdown by ASO targeting the same lncRNA show concordant knockdown profiles, Related to Figure 4. **A**, Diagram showing the method of reciprocal enrichment test, by selecting top and bottom ranked genes from the KD profile and testing for enrichment in another KD profile. **B**, Enrichment tests include forward and reverse analyses and each geneset was tested against all KD profiles from other lncRNA targets to generate sample-based background. **C**, Criteria of calling a hit for concordant ASO-pair. **D**, The numbers of concordant ASO-pairs defined by the criteria shown in (**c**) where all the geneset sizes shown significantly higher number of hits than the different-target-ASO-pair background (empirical *p*-value < 0.05). **E**, An example of concordant pairs defined by n125. **F**, Selection of cutoffs for significant differentially active features by Jaccard index. Under different cutoffs, the Jaccard indexes reflecting the degree of overlap in were taken for the same-target-ASO-pairs (n=293) and the different-target-ASO-pairs (10,000 permutation for 293 pairs). Same-target is significantly higher than different-target in all tested cutoffs (empirical *p*-value < 0.05). Ratios (line plot) represent the mean of same-target-ASO-pairs divided by the mean of different-targets-ASO-pairs. **G**, Jaccard index showing the similarity of isoform target of the 2 ASOs from the same-target-ASO-pairs (n=111) derived from the 36 lncRNAs with concordant molecular phenotype. Upper panel shows the Jaccard index calculated by number of transcript isoforms. Low panel shows the Jaccard index weighted by

**Figure S4.**
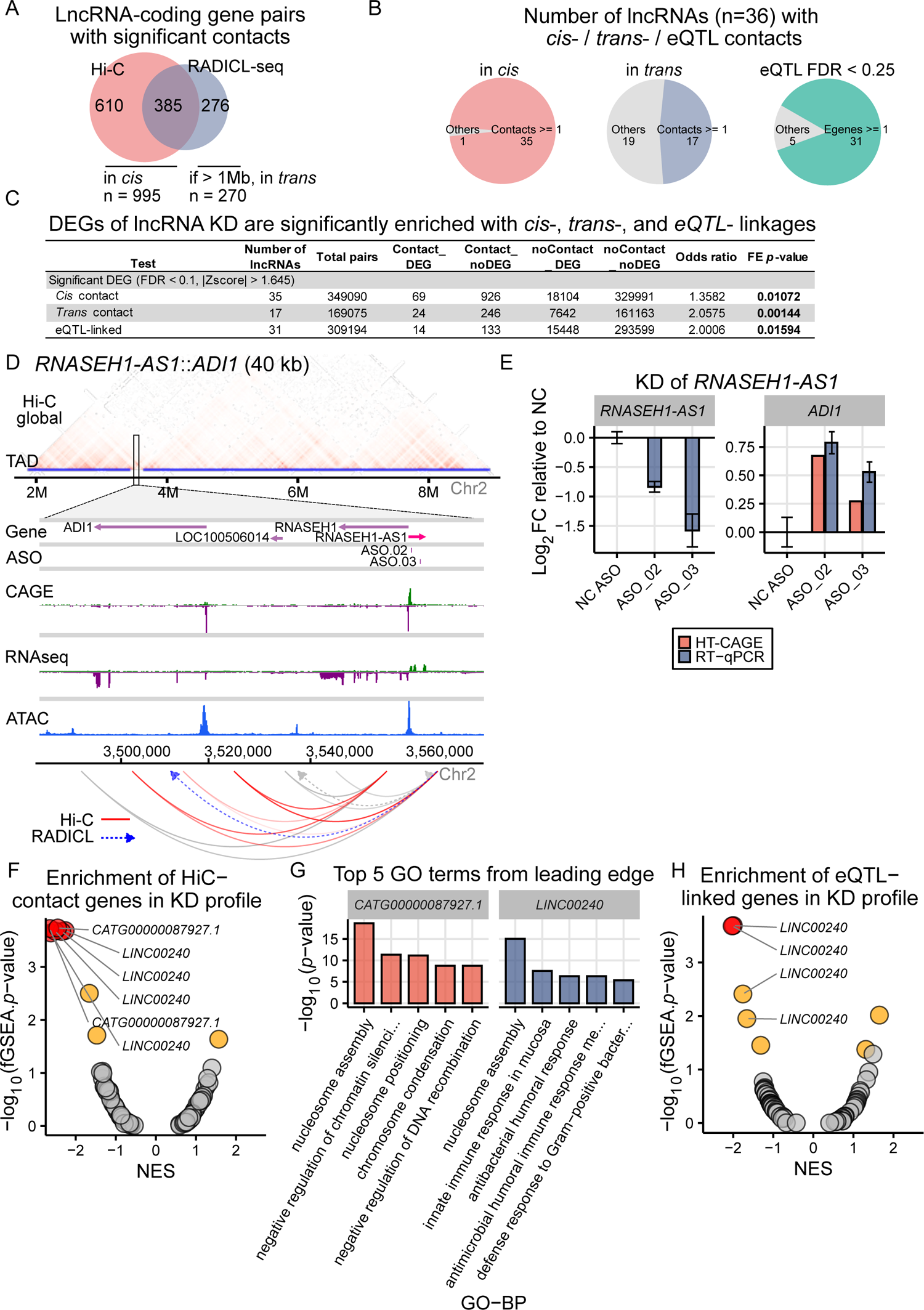
Potential primary targets of lncRNAs identified by knockdown profiles and chromatin interaction, Related to Figure 5. **A**, Venn diagram showing the overlap of Hi-C and RADICL-seq identified lncRNA-protein coding gene pairs. Number of lncRNA-protein coding gene pairs that showed significant interaction (FDR < 0.05) from Hi-C (10 kb resolution) and RADICL-seq (25 kb resolution) data of native iPS cells. Only protein coding genes detectable in HT-CAGE are included. Any pairs with Hi-C contacts are defined as *cis* interaction while pairs with only RADICL-seq contacts and with a minimum genomic distance great than 1 Mb are defined as *trans* interaction. **B**, Number of concordant lncRNAs (n = 36) showing *cis* interaction, *trans* interaction or eQTL-linkage with other protein coding genes. **C**, Fisher’s exact test for the enrichment of *cis*-, *trans*- and eQTL-linkages with significant differentially expressed genes upon knockdown. Only lncRNAs with at least one contact are included while all protein coding genes of selected lncRNAs are used in the analyses. **D**, Another example of Hi-C mediated regulation. Hi-C contacts and TAD regions were defined at the resolutions of 10 kb and 50 kb, respectively. **E**, *RNASEH1-AS1* KD led to up-regulation of *ADI1* from the HT-CAGE result and an independent KD followed by RT-qPCR. **F**, Enrichment of the Hi-C contact gene sets were tested against the KD profiles for each of the 35 lncRNAs (86 ASOs). Red, FDR < 0.05; yellow, *p*-value < 0.05. **G**, TopGo analysis on the leading edge genes for Hi-C contacts enrichment in KD profile showing the top 5 biological process GO terms. **H**, Enrichment of the eQTL linked gene sets were tested against the KD profiles for 31 lncRNAs (77 ASOs).

**Figure S5.**
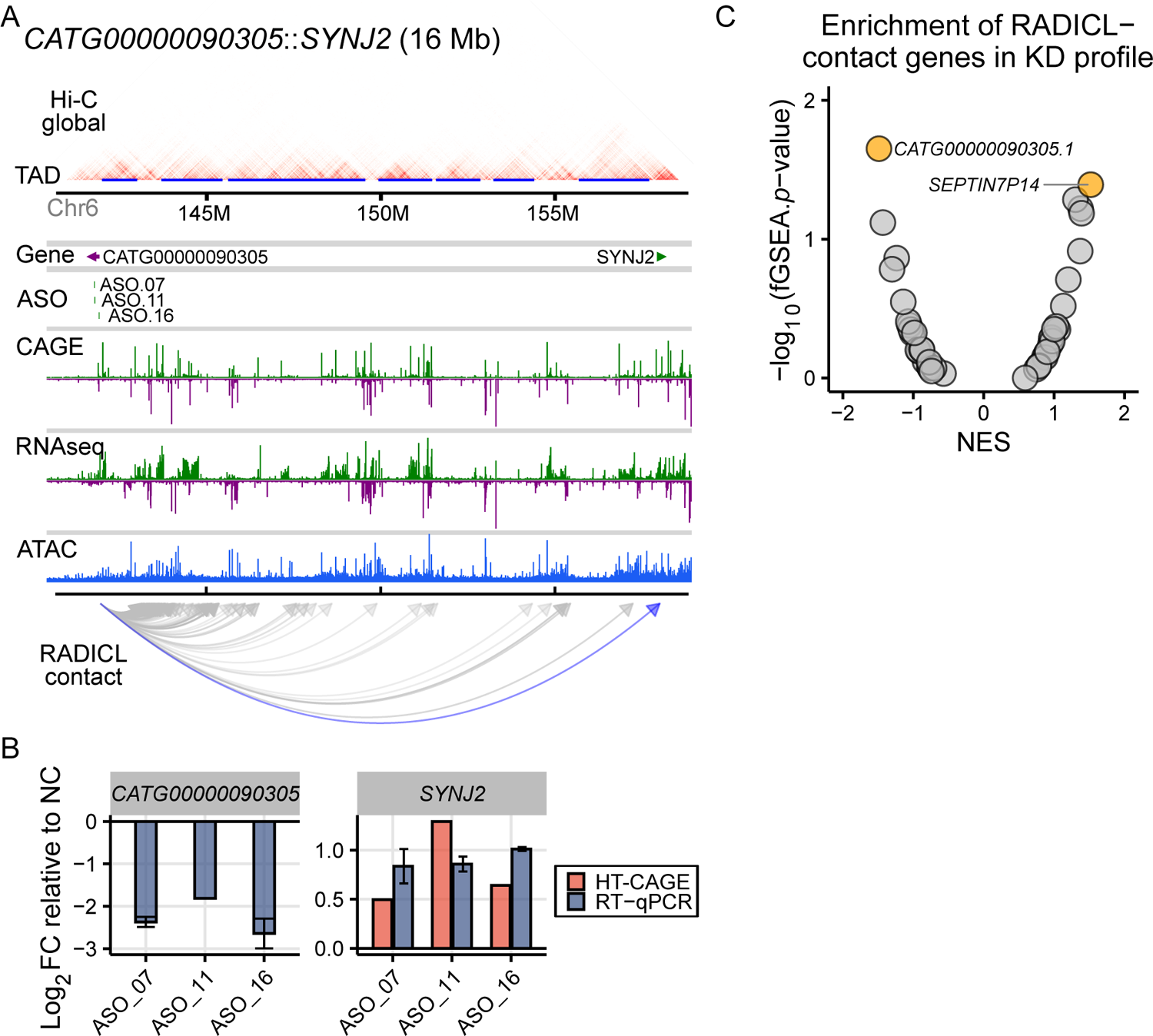
Potential primary targets of lncRNAs identified by knockdown profiles and RADICL-seq, Related to Figure 5. **A**, Genomic view of the RADICL-interaction between *CATG00000090305* and *SYNJ2*. **B**, Up-regulation of *SYNJ2* after the KD of *CATG0000090305* was shown in HT-CAGE and was validated by RT-qPCR. **C**, Enrichment analysis to of the RADICL-seq contact gene sets in each of the 17 lncRNAs (43 ASOs) KD profiles.

**Figure S6.**
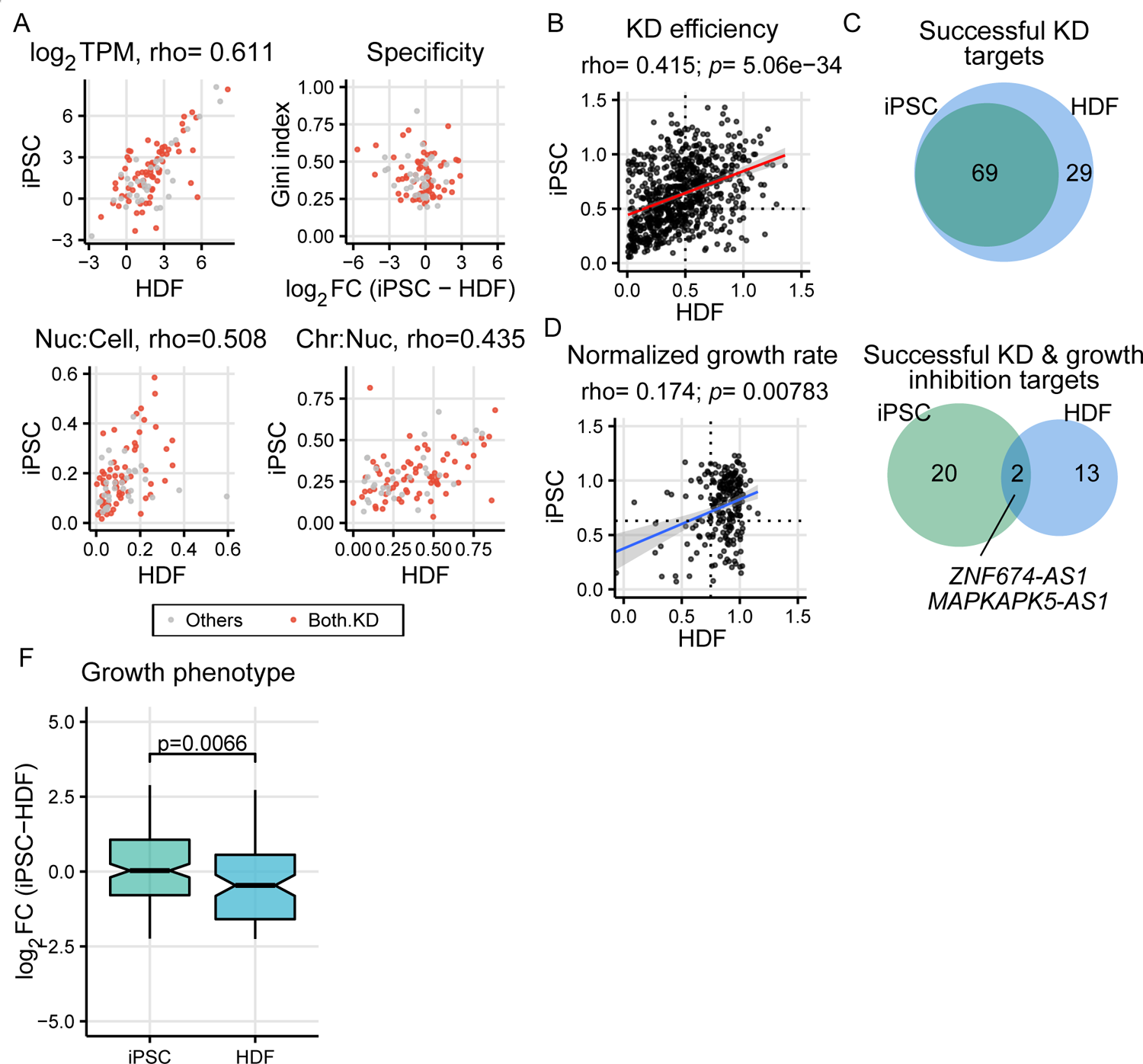
Cell type specificity of lncRNAs between HDF and iPS cells, Related to Figure 7. **A**, Comparison on the expression level, subcellular localization ratio and cell type specificity between HDF (2) and iPS cells of the 102 assayed lncRNAs. **B**, Correlation of KD efficiency at ASO-level. **C**, Venn diagram showing the overlap successful KD at target-level. **D**, Correlation of normalized growth rate of all the effective ASOs. **E**, Venn diagram showing the overlap lncRNAs with growth phenotype at target-level. **F**, Log_2_FC of transcriptomes between HDF and iPS cells was obtained from CAGE followed by edgeR analysis. Log_2_FC of 20 lncRNA targets showing growth phenotypes in iPS cells and 13 targets showing growth phenotype in HDF were compared by Wilcoxon test.

**Figure S7.**
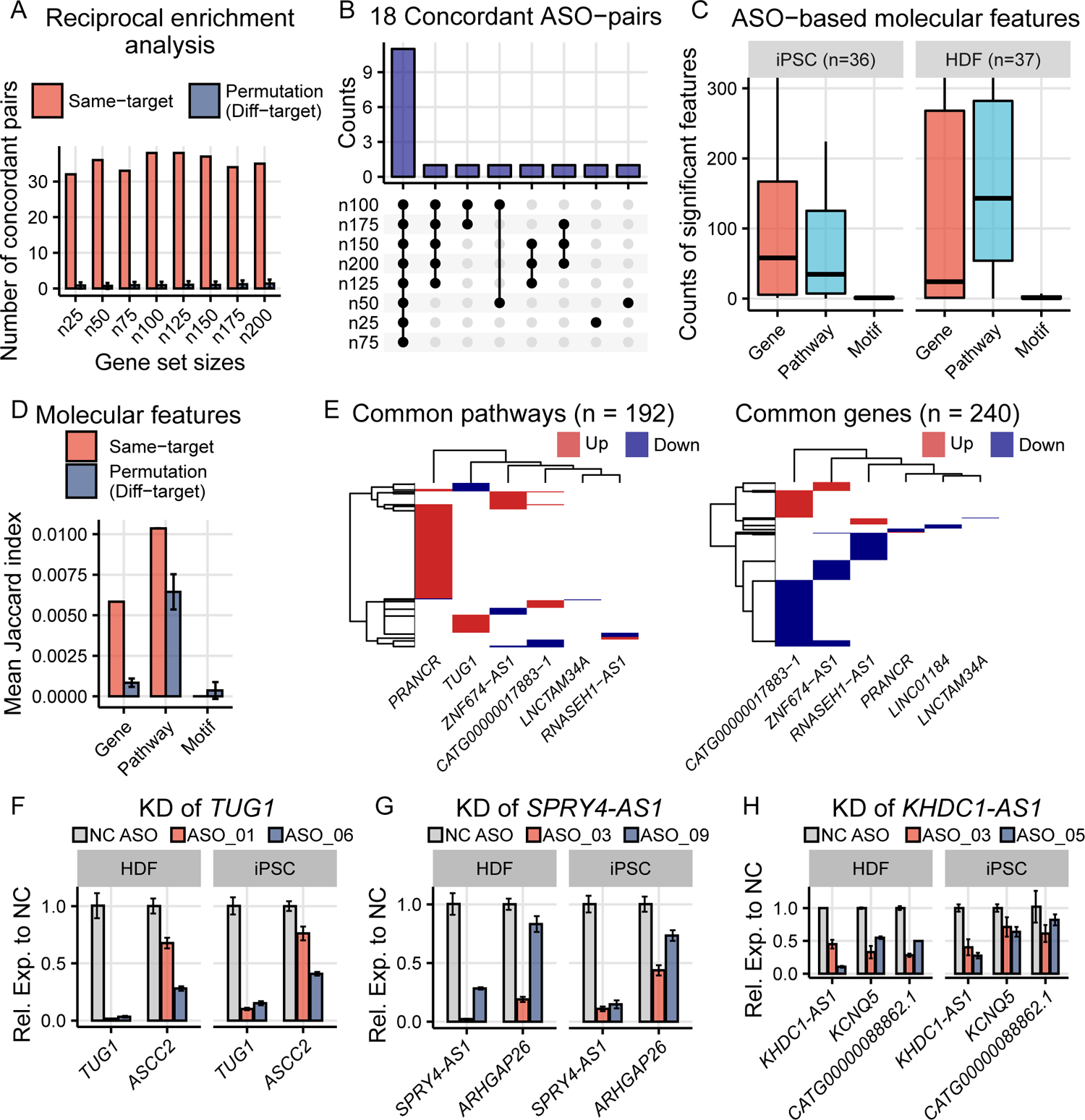
Consistent molecular phenotype of lncRNAs in HDF and iPS cells, Related to Figure 7. **A**, Number of concordant ASO-pairs from the same-target and from the different-target permutation in all the tested geneset sizes. **B**, Plot showing the number of concordant ASO-pairs from different geneset sizes. **C**, Boxplot showing the number of significant differential active molecular features (ASOs from the 12 common lncRNAs) where the lncRNA target is excluded from the count of genes. Significant criteria: iPS-gene, FDR < 0.1, abs(across-sample Z-score) > 1.645; iPS-pathway, FDR<0.05,, abs(across-sample Z-score) > 1.645; iPS-motif, FDR<0.1, abs(across-sample Z-score) > 1.645; HDF-gene, FDR < 0.01, abs(across-sample Z-score) > 1.645, abs(log_2_FC) > 0.5; HDF-pathway, FDR<0.05, abs(across-sample Z-score) > 1.645, abs(NES) > 1; HDF-motif, FDR<0.05, abs(across-sample Z-score) > 1.645. **D**, Using overlap significant differential active features to determine concordance of the same-target-ASO-pairs and compare to different-target pairs with 10,000 permutations. Significant criteria were described in **C**. **E**, Heatmaps showing commonly up- and down-regulated pathways and genes in both HDF and iPSC upon knockdown. The representative ASO-pair used for each lncRNA was shown in **Table S6**. **F&G,** Knockdown of *TUG1* and *SPRY4-AS1* shows regulation on their *cis*-regulated genes in both HDF and iPS cells. **H**, Knockdown of *KHDC1-AS1* resulted in similarly *cis*-regulation on *KCNQ5* and *CATG00000088862.1* in both HDF and iPS cells.

## Notes

https://fantom.gsc.riken.jp/6/suppl/Yip_et_al_2022/

### Competing Interest Statement

The authors have declared no competing interest.

### Summary of Updates

The downstream analyses of the manuscript is restricted to the lncRNAs with significant molecular phenotypes.

